# Uncovering the contributions of charge regulation to the stability of single alpha helices

**DOI:** 10.1101/2022.09.28.509894

**Authors:** Martin J. Fossat, Ammon E. Posey, Rohit V. Pappu

**Affiliations:** Department of Biomedical Engineering and the Center for Biomolecular Condensates, James McKelvey School of Engineering, Washington University in St. Louis, St. Louis, MO 63130

**Author notes:** M.J.F. and A.E.P contributed equally to this work.

## Abstract

The single alpha helix (SAH) is a recurring motif in biology. The consensus sequence has a di-block architecture that includes repeats of four consecutive glutamate residues followed by four consecutive lysine residues. Measurements show that the overall helicity of sequences with consensus E_4_K_4_ repeats is insensitive to a wide range of pH values. Here, we use the recently introduced *q*-canonical ensemble, which allows us to decouple measurements of charge state and conformation, to explain the observed insensitivity of SAH helicity to pH. We couple the outputs from separate measurements of charge and conformation with atomistic simulations to derive residue-specific quantifications of preferences for being in an alpha helix and for the ionizable residues to be charged vs. uncharged. We find a clear preference for accommodating uncharged Glu residues within internal positions of SAH-forming sequences. The stabilities of alpha helical conformations increase with the number of E_4_K_4_ repeats and so do the numbers of accessible charge states that are compatible with forming conformations of high helical content. There is conformational buffering whereby charge state heterogeneity buffers against large-scale conformational changes thus making the overall helicity insensitive to large changes in pH. Further, the results clearly argue against a single, rod-like alpha helical conformation being the only or even dominant conformation in the ensembles of so-called SAH sequences.

## INTRODUCTION

The alpha helix, first discovered by Pauling, Corey, and Branson ^1^ is a recurring secondary structural motif in proteins ^2, 3^. Considerable effort, spanning multiple decades, has gone into uncovering the physico-chemical principles of helix-coil transitions ^4–7^, the sequence determinants of helical stability ^8–11^, rules for helix capping ^12, 13^, the impact of solution conditions on helical stability ^14–17^, cooperative bundling of helices ^18^, and the ability of many systems to form helical structures through coupled folding and binding reactions ^14, 19, 20^. Ionizable residues feature prominently in many helix-forming sequences ^19, 21–23^. Among the roles attributed to ionizable residues is their ability to improve solubility ^11^, their contributions to helical stability through salt bridges ^21, 23^, their preferential accumulation at specific capping positions ^13^, and their ability to contribute to helical stability through pH-dependent uptake or release of protons ^24–28^, which we refer to as *charge regulation* ^29^.

Alpha helices within globular proteins are typically 10-15 residues long ^30^. However, many exceptions exist and the most notable are sequences that form the so-called Single Alpha Helix (SAH) motif. The lengths of such sequences can range from 25 to 200 residues ^31, 32^ (**Figure S1**). Many, although not all SAH formers feature a consensus di-block architecture comprising E_4_K_4_ repeats ^31, 33–35^. Each of the consensus repeats feature four glutamic acid residues followed by four lysine residues. **Figure 1** summarizes the diverse set of functions associated with distinct variants of SAHs, all featuring E_4_K_4_ repeats or variants thereof. In many systems, SAHs are thought to contribute as semiflexible rod-like elements that tether distinct functional domains. Mechanotransduction, mechanical sensing, and force generation are among the numerous functions associated with SAH forming sequences ^36–38^. Further, since their discovery, SAH sequences of different lengths have been used as molecular rulers for measurements of intramolecular distances in cellular systems ^39^.

**Figure 1:**
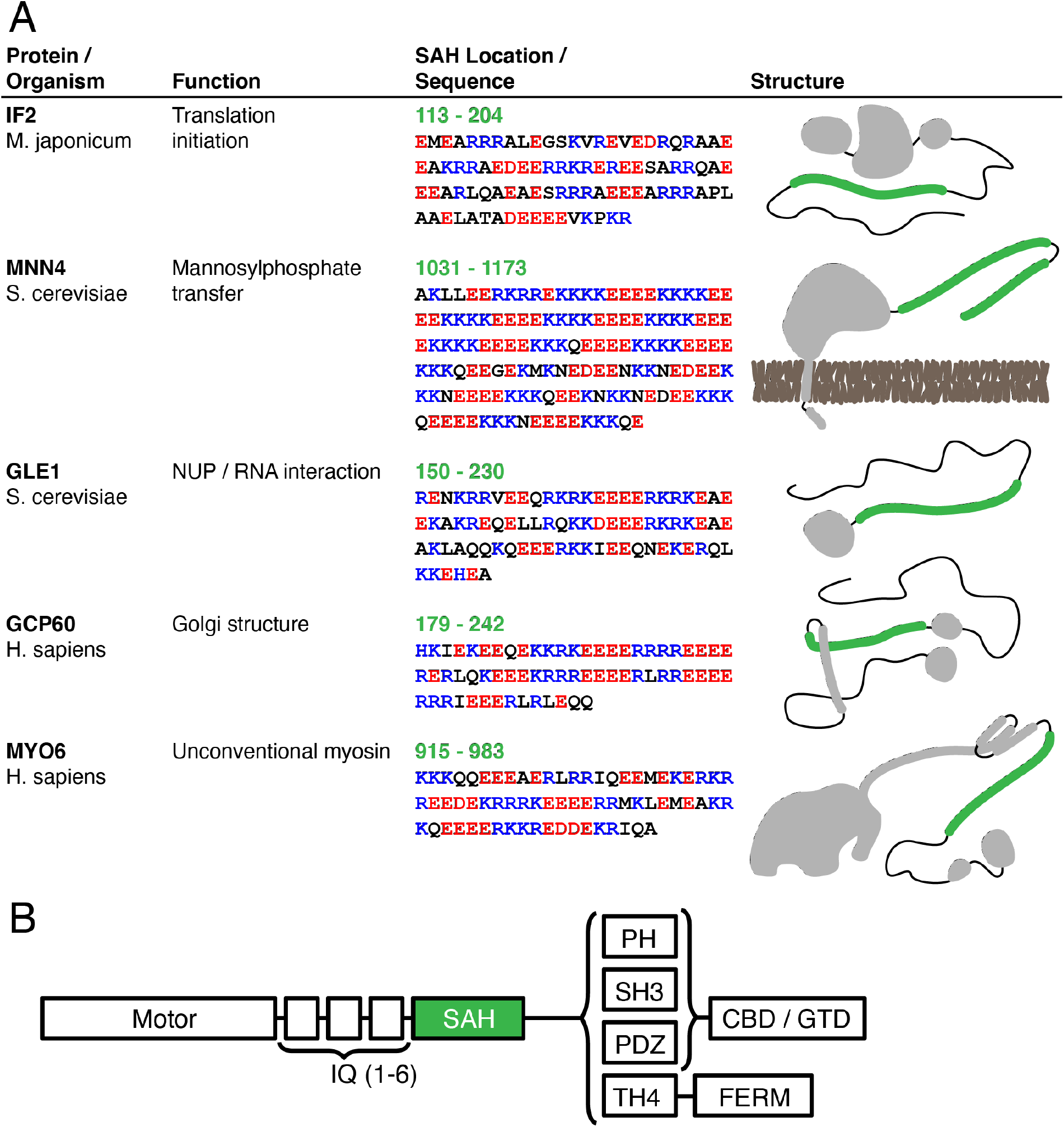
Examples of naturally occurring sequences that are thought to form SAHs. The sequences are from three distinct protein families. (A) The protein name and organism. protein function, SAH amino acid sequence and structural schematics are shown for five representative SAH-containing proteins. SAH regions are colored green in the structural schematics (column 4). For clarity, in column 3, the acidic residues are shown in red, the basic residues in blue, and the constitutively uncharged residues are shown in black. (B) An example consensus domain architecture is depicted for the “unconventional myosin” family proteins that commonly feature SAHs (indicated in green). The domains that follow the SAH are variable, as indicated. The domain architecture schematic was adapted from the work of Fili and Toseland^47^ and Li et al.^48^.

The apparent stability of SAH sequences and their ability to form alpha-helical rod-like conformations have been attributed to *i* to *i*+4 salt bridges between negatively charged Glu residues at position *i* and Lys residues at position *i*+4 ^37, 40^. Two observations suggest that there are likely to be complex contributions to determinants of SAH stability. First, in measurements reported to date, the helicities of SAHs show remarkable insensitivity to changes in pH ^33, 40^. This is noteworthy because the intrinsic pK_a_ values of Arg, Lys, Glu, and Asp are 13.6, 10.7, 4.88, and 4.76, respectively ^41^. Clearly, Glu and Lys are not particularly strong acids or bases, respectively^42^. Therefore, it appears that charge regulation effects are likely contributors to the insensitivity of helical contents as a function of pH. Here, charge regulation refers to the alterations of charge states of ionizable residues through spontaneous binding or release of protons. This hypothesis is bolstered by results from the work of Marqusee and Baldwin ^21^ who showed that the positioning of ion pairs has a modest effect on alpha helicity in alanine-based systems. They also showed that the alpha helicity in model alanine-based peptides that feature complementary ion pairs between Arg / Lys and Glu residues can be higher at extremes of pH when compared to pH 7.5. Under these conditions, the peptides have a net charge as opposed to being electroneutral. Additionally, Doig and coworkers have shown that the positioning of phosphoserine in alanine-based systems has a profound effect on helical stability ^43, 44^. These observations were linked to the position-dependence of the charge state of the phosphoserine and lysine residues within the sequence. Together, these studies suggest that charge regulation is likely to be an important contributor to the stability of SAH forming sequences. Further support for the putative role of charge regulation comes from a second observation pertaining to the imperfectness of E_4_K_4_ repeats. Although E_4_K_4_ is the consensus motif, one often sees interruptions of this pattern that go beyond the substitution of Lys with Arg. Indeed, Glu and Lys are often substituted with uncharged residues such as Ala, Gln, or Leu (**Figure 1**) ^31^. This can give rise to sequences with a net charge, as naively evaluated by assuming the charge states to be fixed by the pK_a_ values of model compounds. Additionally, while there are occasional substitutions of Glu with Asp, one never observes blocks of Asp residues^45^. The likelihood of observing four consecutive Arg residues is also lower than the likelihood of observing four consecutive Lys residues.

Why might charge regulation have a role to play in helix formation / stabilization? First, Pace and Scholtz measured the intrinsic helical propensities of naturally occurring amino acids and uncharged versions of specific ionizable residues. Averaged over three different sequence contexts, the consensus helical propensities that were derived from their measurements have the following hierarchy: Glu° > Ala > Leu ≈ Arg^+^ > Lys^+^ > Gln > Glu^−^ > Asp° > Asn > Asp^−^. Here, Glu° and Asp° are uncharged versions of Glu and Asp, respectively. Likewise, Arg^+^, Lys^+^, Glu^−^, and Asp^−^ are charged versions of Arg, Lys, Glu, and Asp, respectively. Clearly, the intrinsic helical propensities are highest for Glu°. Second, the free energies of hydration, which are relevant for desolvation penalties associated with helix formation, are different for basic vs. acidic groups, and considerably more favorable for Asp° and Glu° compared to Asp^−^ and Glu^− 41, 46^. This points to the possibility of lowering the desolvation penalty through site-specific or context-dependent proton uptake or release. Third, a strong case has been made for the importance of reducing the linear charge density along to the helical axis ^22^ – a feature that is achievable through charge regulation. And fourth, local electrostatic repulsions within and between like-charged blocks are likely to disrupt or destabilize optimal *i* to *i*+4 salt bridges.

Taken together, the observations summarized above suggest that modulations of charge states of ionizable residues are likely to impact SAH stability through increased intrinsic helical propensities, lowered charge density along the helical axis, and decreased desolvation penalties. Further, selective, position-specific neutralization of ionizable residues could enhance the specificity of helix-stabilizing *i* to *i*+4 salt bridges by avoiding the possibility of alternative salt bridges. To test these hypotheses, we performed systematic investigations of the pH-dependent behaviors of three systems with E_4_K_4_ repeats.

Our investigations were aided by the *q*-canonical ensemble ^29^, which we recently introduced. This allows us to parse measurements based on potentiometric titrations to characterize the ensemble of charge states that are prevalent in solution for a specific pH ^29^. The *q-*canonical ensemble was applied to characterize the ensemble of charge states that peptides with E_4_K_4_ repeats can access as a function of pH ^29^. Here, we adapt this formalism by combining the measurements of charge states with measurements of conformations to go alongside atomistic simulations based on the ABSINTH implicit solvation model and forcefield paradigm ^49^. The combined approach helps us uncover the role of charge regulation as a determinant of the insensitivity of SAH stability to changes in pH. Increases in charge state heterogeneity do not compromise helical stability. Our results suggest that the effects of proton binding and unbinding are likely to add to the contributions of backbone hydrogen bonding, backbone solvation / desolvation, and inter-block salt bridges as determinants of SAH stability.

Our studies focus on three systems of the form Ac-G-(E_4_K_4_)_n_-GW-NH_2,_ where n = 1, 2, and 3, respectively. Here, Ac refers to N-acetyl, and the tryptophan residue was used to enable precise measurements of peptide concentration. As a shorthand, we refer to the three systems as (E_4_K_4_)_1_, (E_4_K_4_)_2_, and (E_4_K_4_)_3_, respectively. First, we present results summarizing the repeat number and pH dependence of helicity as measured by ultraviolet circular dichroism (UV-CD). Then, we summarize the main tenets of the *q-*canonical ensemble, specifically the parsing of the system into microstates and mesostates. This is followed by a summary of the workflow that combines potentiometric titrations, reported previously, with the UV-CD measurements to extract intrinsic helicities of different charge mesostates. Using mesostate-specific helicities as restraints, we performed atomistic simulations based on the ABSINTH implicit solvation model to extract a full description of the pH-dependence of conformations and charge states that are consistent with the experimental data. These results were used to extract insights regarding the pH-dependent preferences for charge states of individual ionizable residues. For helical conformations, we uncovered clear, repeat number dependent downshifts in the apparent pK_a_ values of internal Glu residues and significant, position-specific apparent pK_a_ values for Lys residues. Further, we observed increased charge state heterogeneity with increasing number of repeats. This prevails despite negligible changes to the overall helicity. Our results suggest that at a given pH, the overall ensemble, especially for longer sequences, is best described as being predominantly helical while accommodating significant charge state heterogeneity. However, our results highlight the fact that the single, continuous helix is not a prominent feature of the conformational ensemble, especially for the finite-sized systems we study in this work.

## RESULTS

### (E_4_K_4_)_n_ systems show n-dependent and pH-independent preferences for overall helicity

We performed UV-CD measurements for each of the three (E_4_K_4_)_n_ systems, where n = 1, 2, and 3 (**Figure S2**). The measurements were performed in a background solution of 10 mM KCl and 5 mM HCl at a temperature of 20°C and titrated across pH values ranging from 2 – 12 using KOH. The overall helicity, measured using UV-CD, and plotted as the fractional helicity, is summarized in **Figure 2**. We observed the following trends in the data: For a given pH, the fractional helicity increases with the number of E_4_K_4_ repeats. For each construct, there is a pH window where the fractional helicity changes minimally. The size of this window increases with the number of repeats. In the pH window where the helicity varies minimally, the fractional helicity values are ≈0.17 for (E_4_K_4_)_1_, ≈0.5 for (E_4_K_4_)_2_, and ≈0.7 for (E_4_K_4_)_3_. These helicities are concordant with values reported in the literature ^34^. The key result is the insensitivity of overall helicity to significant changes in pH.

**Figure 2:**
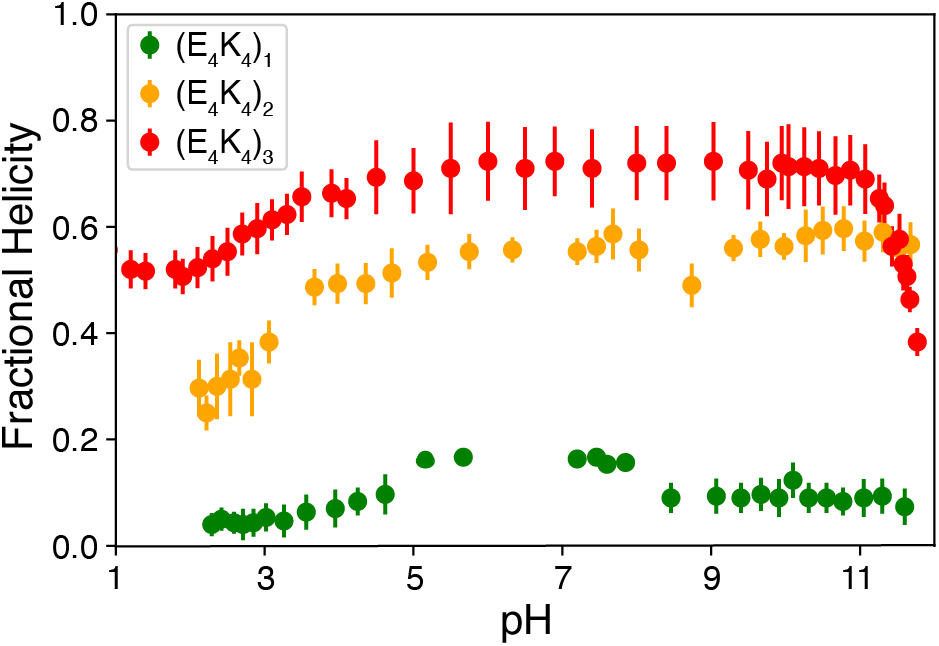
Fractional helicity derived from the UV-CD data. The full spectra are shown in **Figure S2**. Data points and error bars respectively represent the mean fractional helicity from deconvolution and the standard deviation of deconvolution using three different basis sets, as described in the Methods section

### Potentiometric titrations enable the measurements of charge states of (E_4_K_4_)_n_ peptides

In a recent report, we published results from potentiometric titrations of the three (E_4_K_4_)_n_ peptides studied in this work ^29^. The salient results from these titrations are summarized in **Figure S3**. Potentiometry yields estimates of the net charge as a function of pH. We used the formalism of the *q-*canonical ensemble, introduced previously ^29^, to extract information regarding the distinct charge states that contribute to the titrations as a function of pH. The overall approach is summarized below using the (E_4_K_4_)_1_ peptide as an illustrative system.

There are eight ionizable residues for the (E_4_K_4_)_1_ peptide. Each Glu residue can be charged or uncharged. These are denoted as E and e, respectively. Likewise, each Lys residue can be protonated (K) or deprotonated (k). In all, for (E_4_K_4_)_1_ there is a theoretical maximum of 2^8^ or 256 charge microstates, where each microstate is denoted as a specific sequence *viz*., eEEEKKKK, EeEEKKKK, etc. Of these, several microstates are forbidden since Lys and Glu residues are unlikely to be simultaneously uncharged. Accordingly, for (E_4_K_4_)_1_, the number of relevant charge microstates reduces to 32 as was shown previously ^29^. Next, the charge microstates are grouped into mesostates. Each mesostate encompasses all microstates that have the same net charge. Mesostates are given an X_y_ designation, where X refers to the net charge associated with each of the microstates within a mesostate, and y refers to the number of thermodynamically accessible microstates within a mesostate (**Figure S4**).

The pH-dependence of the net charge is used to extract the probability that a specific mesostate will be populated at a given pH. Previous work showed that for (E_4_K_4_)_1_, the mesostate namely 0_1_ dominates in the pH range between 7 and 9 ^29^. However, the data also indicate a downshift vs. an upshift in the pK_a_ values for the Glu and the Lys, respectively. For the longer peptides (E_4_K_4_)_2_ and (E_4_K_4_)_3_, there are at least two mesostates of net charge ±1 that contribute to the measured charge profiles at every pH, even if the contributions are small (**Figure S5**). The pH dependent populations of mesostates can be used to quantify the numbers of mesostates that contribute to 95% vs. 99% of the overall population as a function of pH, as shown in **Figure S6**.

The overall picture that emerges is of an ensemble of distinct charge microstates. The number of thermodynamically relevant microstates / mesostates is set by the pH. The 0_1_ mesostate becomes less important outside the pH window of 7.0 – 7.5. Accordingly, the question is if there are mesostates with a net charge that also have high intrinsic helicities. The existence of such mesostates would be one way to explain the robust pH-independence of helicity of (E_4_K_4_) repeats.

### Results from potentiometric titrations and pH-dependent UV-CD measurements can be combined to quantify the intrinsic helicities of mesostates

The pH-dependent mesostate probabilities *p_q_*(pH), extracted from potentiometric titrations, can be used in conjunction with the pH-dependent estimates of overall helicity, f_α_(pH) plotted in **Figure 2**, to extract intrinsic helicities, f_*α,q*_, for each of the mesostates characterized by a net charge *q*. For this, we use the formalism summarized in Equation (1) in conjunction with a Monte Carlo fitting procedure (see details in the methods section).

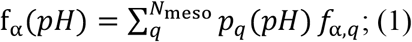

Here, f_α_(pH) and *p_q_*(pH) are inferred using data from UV-CD measurements and potentiometric titrations, respectively. These are then used in a Monte Carlo fitting procedure to extract f_*α,q*_. The workflow is summarized in panels (A) and (B) of **Figure 3**.

**Figure 3:**
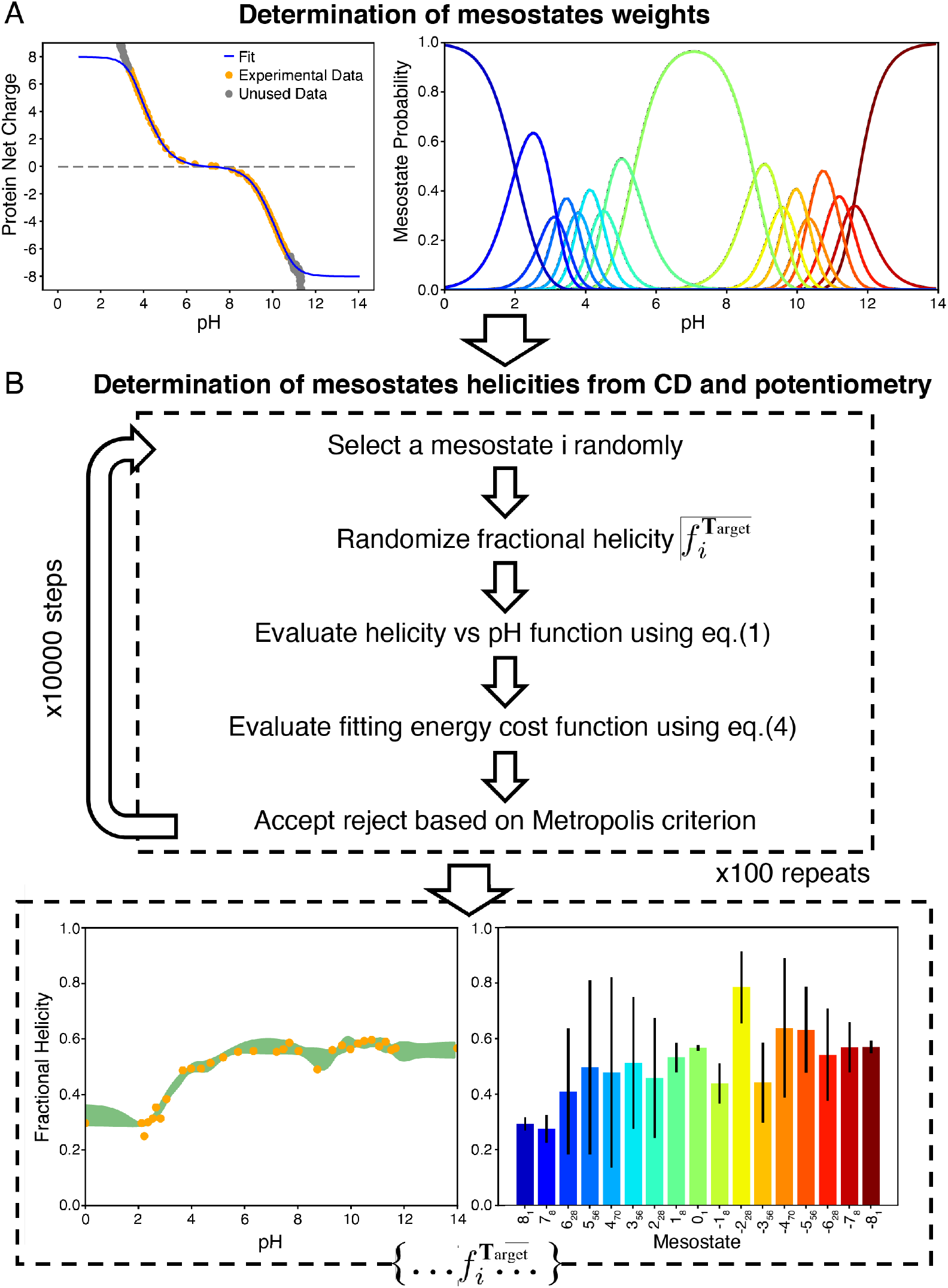
Workflow for deploying the full *q*-canonical ensemble using potentiometric titrations, UV-CD spectroscopy, and atomistic simulations. (A) Summary of the fitting methods used to extract mesostate probabilities as a function of pH from potentiometric titrations. The approach is that of Fossat et al. ^29^. (B) Summary of the procedure to determine the intrinsic fractional helicities of mesostates. Here, we use the mesostate probabilities derived from the workflow shown in panel (A). This is combined with pH-dependent CD measurements.

The values of f_*α,q*_, extracted from the fitting procedure, are summarized in **Figure 4A-4C**. Further, as shown using the solid curves in **Figure 4D**, the values obtained for f_*α,q*_ can be used to obtain fits to the measured fractional helicities as a function of pH. The key takeaway is that there are several mesostates, especially for the (E_4_K_4_)_2_ and (E_4_K_4_)_3_ systems whose intrinsic helicities are equivalent to or greater than that of the 0_1_ mesostate. All these mesostates have a net charge. The intrinsic helicities of some of the mesostates with a net charge including those with a net charge of +12, +9, +2, −11 and −12 are equivalent to or higher than the intrinsic helicity of the 0_1_ mesostate for the (E_4_K_4_)_3_ system.

**Figure 4:**
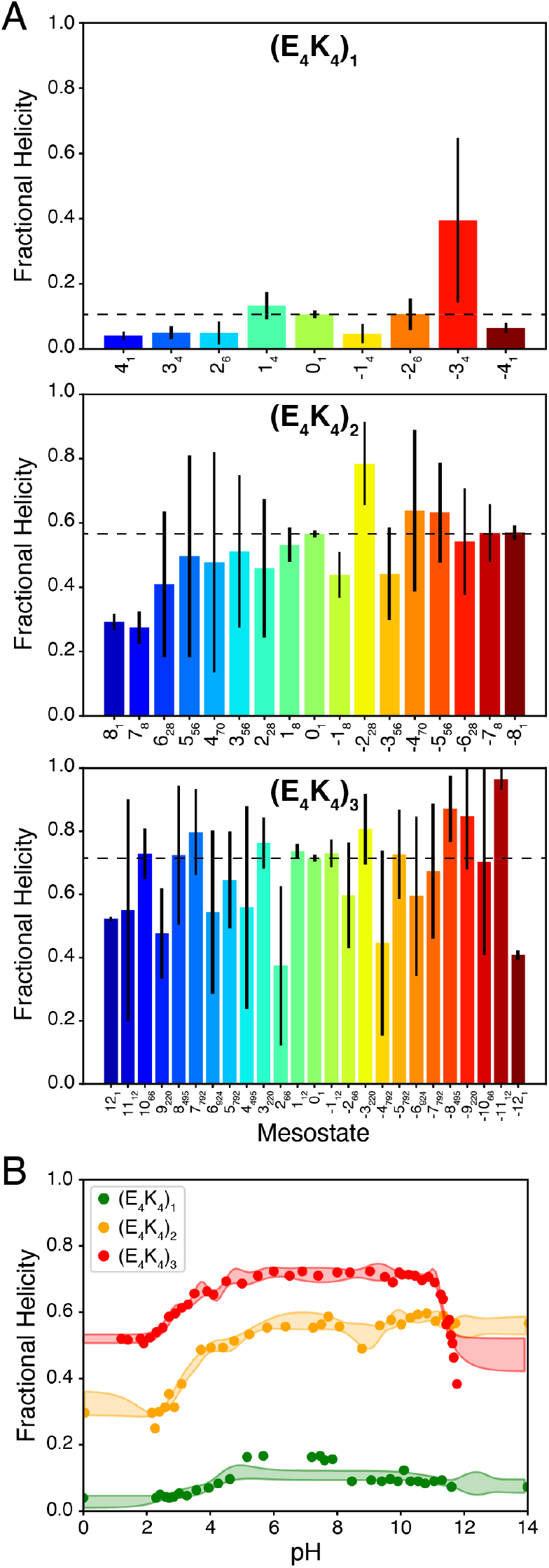
Bar plots summarizing the intrinsic helicities, shown as fractions that lie between 0 and 1. **(A)** Data are shown for the relevant mesostates for each of the (E_4_K_4_)_n_ peptides. In the three sub-panels that make up (A), the dashed line quantifies the construct specific intrinsic helicity of the 0_1_ mesostate. The intrinsic helicities were extracted using a combination of mesostate weights from potentiometric measurements and from the CD data. (B) The figure shows mean fractional helicities determined from deconvolution of the CD data. The fitting envelope corresponds to the standard deviation of the best fits from each of the 100 fits as a function of pH. These data were jointly analyzed as summarized in the main text and panel (B) of Figure 3. The analysis is based on the use of Equation 1.

The potentiometric titrations help identify the mesostates that are relevant for a coarse-grained description of the overall *q-*canonical ensemble. By combining information from potentiometric titrations with pH-dependent UV-CD measurements, we quantify the intrinsic helicities of mesostates. To obtain information regarding the microstates, specifically the types of charge microstates and the conformations of microstates that contribute to the overall ensemble, we turn to simulations as summarized in **Figure 5**.

**Figure 5:**
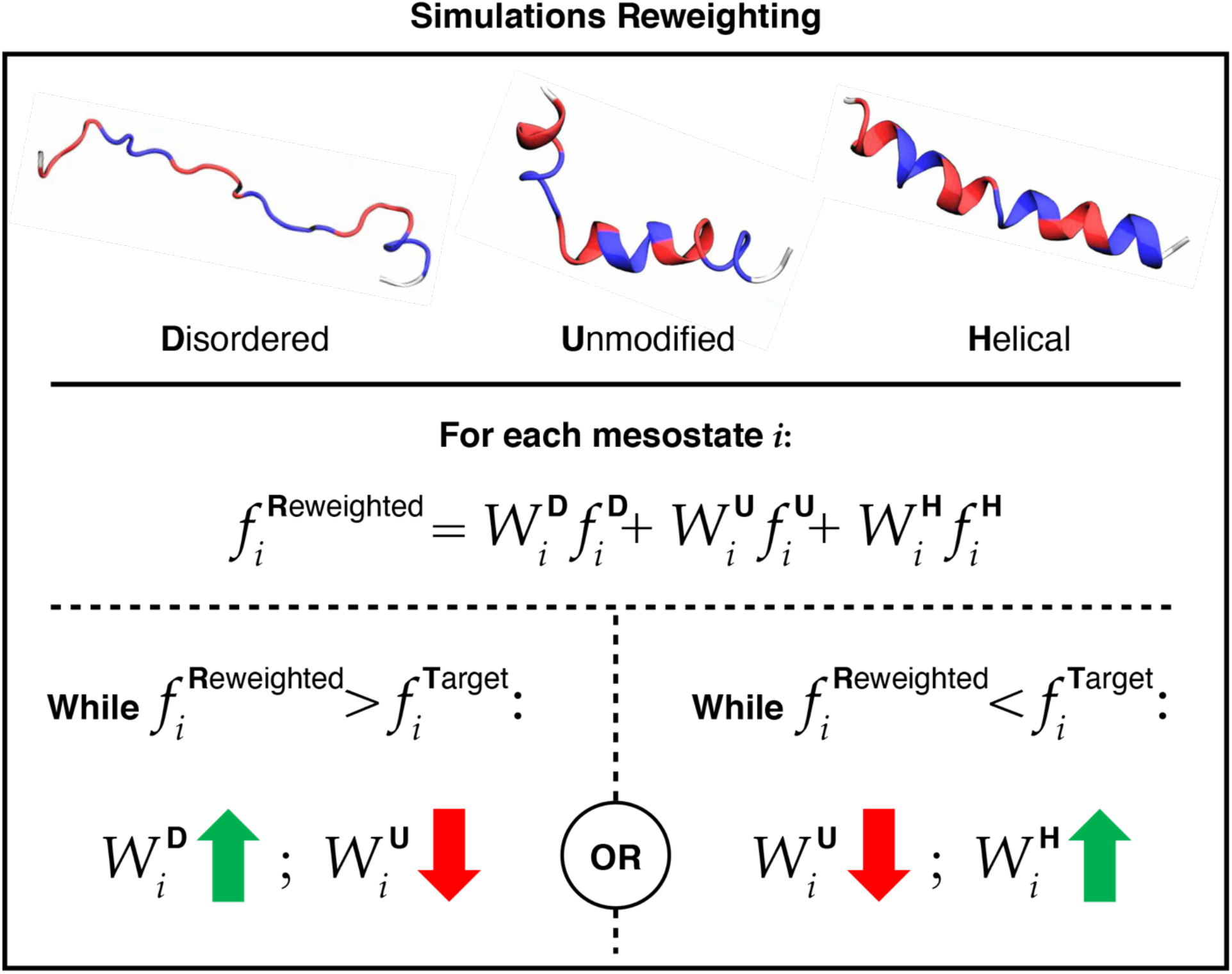
Summary of the procedure used to mix conformational ensembles from three distinct sets of simulations and reweight the overall ensemble to gain joint agreement with potentiometric titrations and UV-CD data. Here, *f*_i_^D^, *f*_i_^U^, and *f*_i_^H^ are the contributions to fractional helicities of mesostates from the disordered ensemble D, the unbiased or unmodified ensemble U, and the helical ensemble H. Likewise, *W*_i_^D^, *W*_i_^U^ and *W*_i_^H^ are the weights of the mixing function, which are refined through iteration. The converged values of these weights reflect the contributions from the three ensembles that yield congruence with the experimental data for fractional helicities of mesostates.

### Atomistic simulations identify the microstates that contribute to the experimentally observed helicity profiles as a function of pH

The simulations use atomic-level descriptions of polypeptides and solution ions based on an all-atom forcefield. The aqueous solvent is modeled implicitly using the ABSINTH model and forcefield paradigm ^49^. For each system, a series of simulations were performed for each of the mesostates using the *q*-canonical Monte Carlo (*q*MC) sampling methodology ^50^. Further, we used the combination of inferred intrinsic helicities of mesostates and their populations as a function of pH as inputs for reweighting the simulations for each of the mesostates. The reweighting procedure is summarized in **Figure 5**.

For reweighting, we mix conformations from three distinct ensembles *viz*., conformations obtained from a highly disordered ensemble obtained by setting the simulation temperature to 350 K, a second set corresponding to unbiased simulations performed at a simulation temperature of 240 K, and the third set corresponding to the most helical ensemble. The helical ensemble is obtained by imposing harmonic constraints on the backbone, thus maintaining an overall helical conformation. The temperature of 240 K for the unbiased simulations was chosen using simulations initiated within the 0_1_ mesostate and identifying the temperature at which the helicity of this mesostate equals the helicity extracted from the combination of CD and potentiometry. It is worth noting that simulation temperatures and the real temperatures in experiments do not have a one-to-one congruence for classical forcefields and implicit solvation models.

A suitable mixture of the three sets of simulated ensembles is obtained by reweighting on a per mesostate basis as summarized in **Figure 5**. For each mesostate, we assess the overall helicity in each of the three sets. We then compare the mesostate helicities of the unmodified set to the mesostate helicity obtained from experiments. If the helicity is too high for that mesostate compared to the experimentally derived value, we introduce the first set until the overall weighted sum of the two sets is within 0.001 of experiments. Conversely, if the helicity is too low for a mesostate compared to the experimentally derived value, we introduce conformations from the disordered ensemble to converge on the overall reweighted ensemble. The proportion of each ensemble used per mesostate is shown in **Figure S6**.

Using the mixing function shown in **Figure 5**, we derived weights associated with each of the three ensembles for each mesostate. We then set the mesostate free energies to those obtained from potentiometric data, while conserving the properly mixed microstate weights within mesostates. This means that the average free energy of each mesostate is concordant with potentiometrically derived values, and the free energy differences among microstates within mesostates are those derived from simulations, after proper mixing. This allows us to extract microstate weights that are also consistent with experiments thereby allowing us to generate structural insights on a per residue basis. Our use of the three sets of simulated ensembles, including the unbiased simulations allows us to obtain a per residue profile that reflects the physics of the forcefield paradigm used in our simulations.

The salient results from the reweighting are summarized in **Figure 6**. Here, each row shows results for a specific pH, whereas each column shows results for a specific construct. The quantities being plotted are the contributions of distinct mesostates to the position-specific helicity values. The latter were derived from the properly reweighted, pH-dependent ensembles of microstates. Along each column, we note that the heterogeneity of charge states is highest at pH 9.5 and 5.5 and lowest at pH 7.5. Further, going across each row, the contributions from charge state heterogeneity to the overall *q*-canonical ensemble become more pronounced with increasing numbers of E_4_K_4_ repeats. The implication is that as the number of repeats increases, there are more charge states that are compatible with similar degrees of helicity. This is especially true at pH values that are above or below pH 7.5.

**Figure 6:**
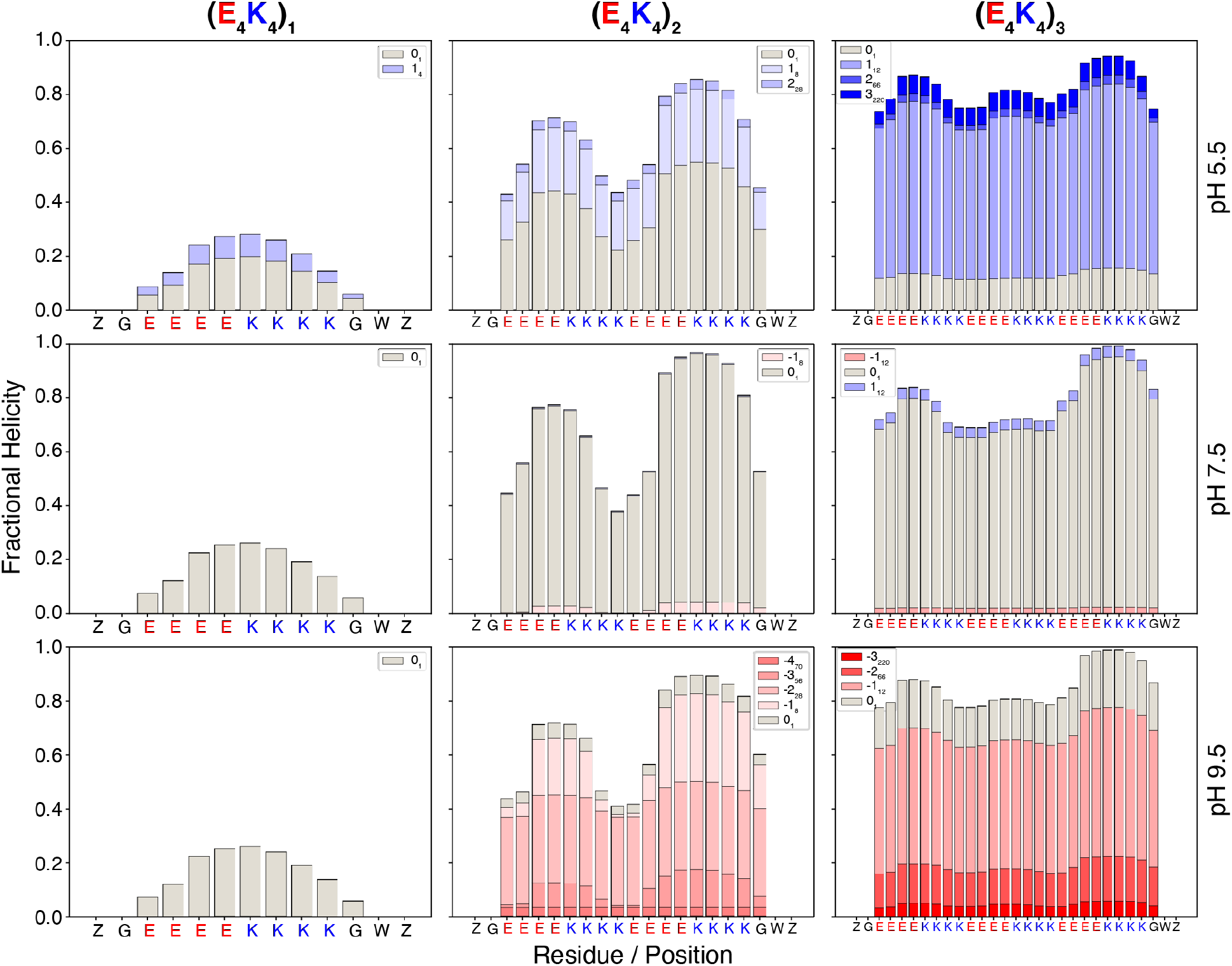
Mesostate-specific contributions to helicity per residue at three different pH values. Each column contains data for a specific construct, and each row contains data for a specific pH. The stacked bars quantify the contributions from specific mesostates to the overall helicity per residue. The insets show the relevant color coding of the mesostates within a panel. Here, Z along the abscissa refers to the N- and C-terminal caps. At pH 5.5, the relevant mesostates for (E_4_K_4_)_3_ have a net charge of +1, +2, and +3. The mesostates with net charge of +1, +2, and +3 have 12, 66, and 220 accessible microstates, respectively and the likelihoods of accessing these microstates are determined by the intra-mesostate microstate probabilities. In terms of their contributions to the overall helicity, the dominant mesostates follow the hierarchy, +1_12_ > +3_220_ ≈ 0_1_ > +2_66_. At pH 7.5, there are minor contributions from the mesostates with charge of ±1 for (E_4_K_4_)_3_, and the hierarchy in terms of contributions to the overall to the overall helicity is 0_1_ >> +1_12_ > −1_12_. Finally, for (E_4_K_4_)_3_ at pH 9.5, the contributions to the overall helicity from mesostates follow the hierarchy, −1_12_ > −2_66_ ≈ 0_1_ > −3_220_. These results illustrate the contributions of charge state heterogeneity to the maintenance of overall helicity. It is worth noting that helicities of Z and G are not zero. Instead, the plots we show reflect our choice of the DSSP algorithm ^3^ and its implementation in mdtraj^51^ for quantifying per-residue helical contents. The algorithm relies on the identification of hydrogen bonds to define whether a structure is in an alpha helical conformation. This relies on the identification of chiral centers and the presence of four consecutive alpha carbon atoms. Accordingly, the two terminal groups namely Z and G, are by the choice of the algorithm, not included in computations of helicity.

### How rod-like are the conformational ensembles at a given pH?

Next, we quantified the distributions of the lengths of helical segments within the reweighted ensembles. The results are shown in **Figure 7** for (E_4_K_4_)_3_ and in **Figure S7** for (E_4_K_4_)_2_. The overall helicity is low for (E_4_K_4_)_1_. Despite a high fractional helicity of ≈0.7 that persists across the pH-dependent ensembles, the perfectly helical rod-like conformation is not the dominant species for (E_4_K_4_)_3_. This is especially relevant, as a cautionary note, given the use of SAH-forming sequences as molecular rulers for intracellular distance measurements ^39^. The overall ensemble is a mixture of partial and fully helical conformations. The four main conformational classes are as follows: In class 1, the N- and C-terminal repeats are alpha helical, whereas the internal repeats are disordered. In class 2, roughly 50-60% of the sequence encompassing the N-terminal portion adopts alpha helical conformations whereas the remaining C-terminal portion is disordered. The converse is true for class 3. Finally, in class 4, conformations correspond to the fully alpha helical, rod-like state (**Figure 6**). The distributions of alpha helical segments of specific lengths share commonalities across the different pH values. There are three distinct peaks corresponding to class 1, class 3, and class 4 conformations, respectively. Class 2 conformations correspond to a trough between two peaks associated with class 1 and class 3 conformations. The trough at an alpha helical segment length of 20 residues points to a clear preference against disrupting the transition between class 3 and class 4 conformations.

**Figure 7:**
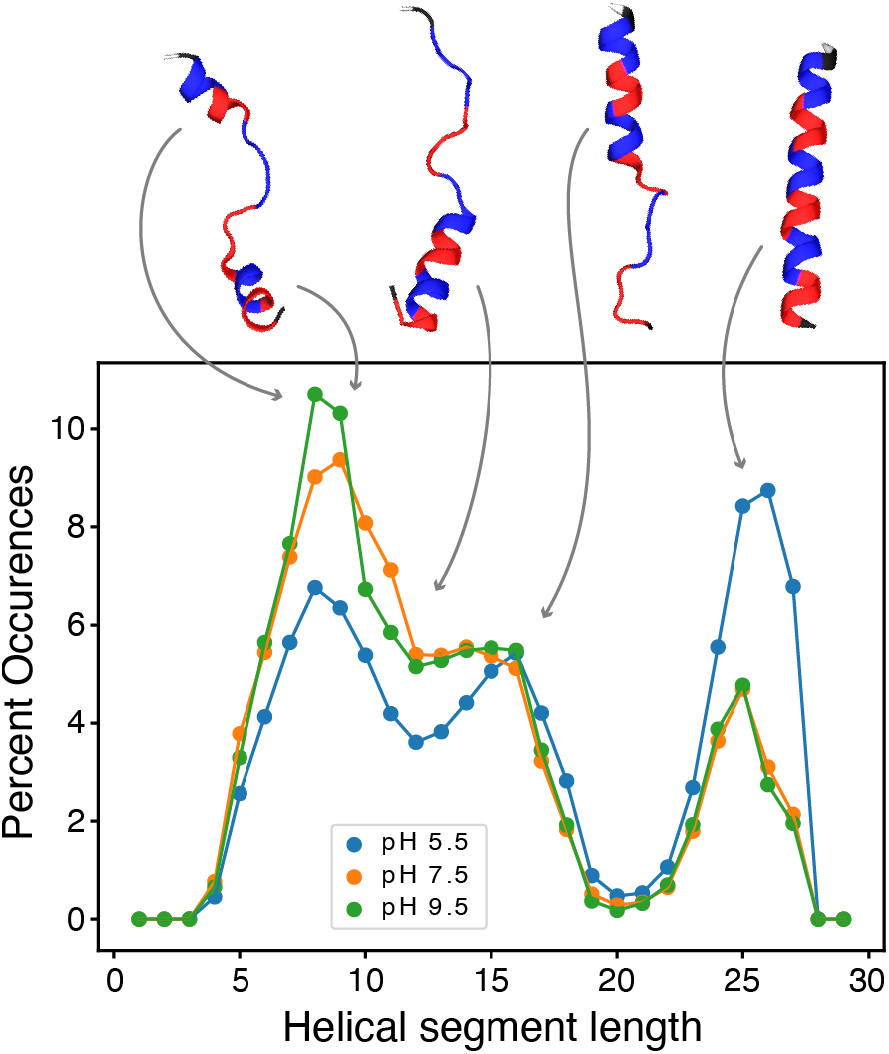
Distributions of the lengths of helical segments extracted from the reweighted ensembles of (E_4_K_4_)_3_ at pH 5.5, 7.5, and 9.5. The top row shows representative conformations as annotations of specific features of the graph. Conformations corresponding to classes 1 – 4 are depicted going from left to right.

### Site specific ionization preferences are context dependent

In **Figure 8** we plot the probability of finding each residue in a charged state. For simplicity of presentation, the data are shown for (E_4_K_2_)_2_. For reference, ionization curves are also shown for the case where we assume fixed charge states based on the pK_a_ values of model compounds. We define the site-specific apparent pK_a_ as the pH at which the likelihood of being charged is 0.5. The top rows of columns A and B of **Figure 8** show insets that zoom into the ionization curves with the dashed lines highlighting the likelihood of 0.5. Overall, the apparent pK_a_ values inferred from the ionization curves suggest that the Glu residues have systematically downshifted pK_a_ values when compared to the model compound value of 4.34 ^52^. The maximal downshift is approximately 0.8 of a pH unit.

**Figure 8:**
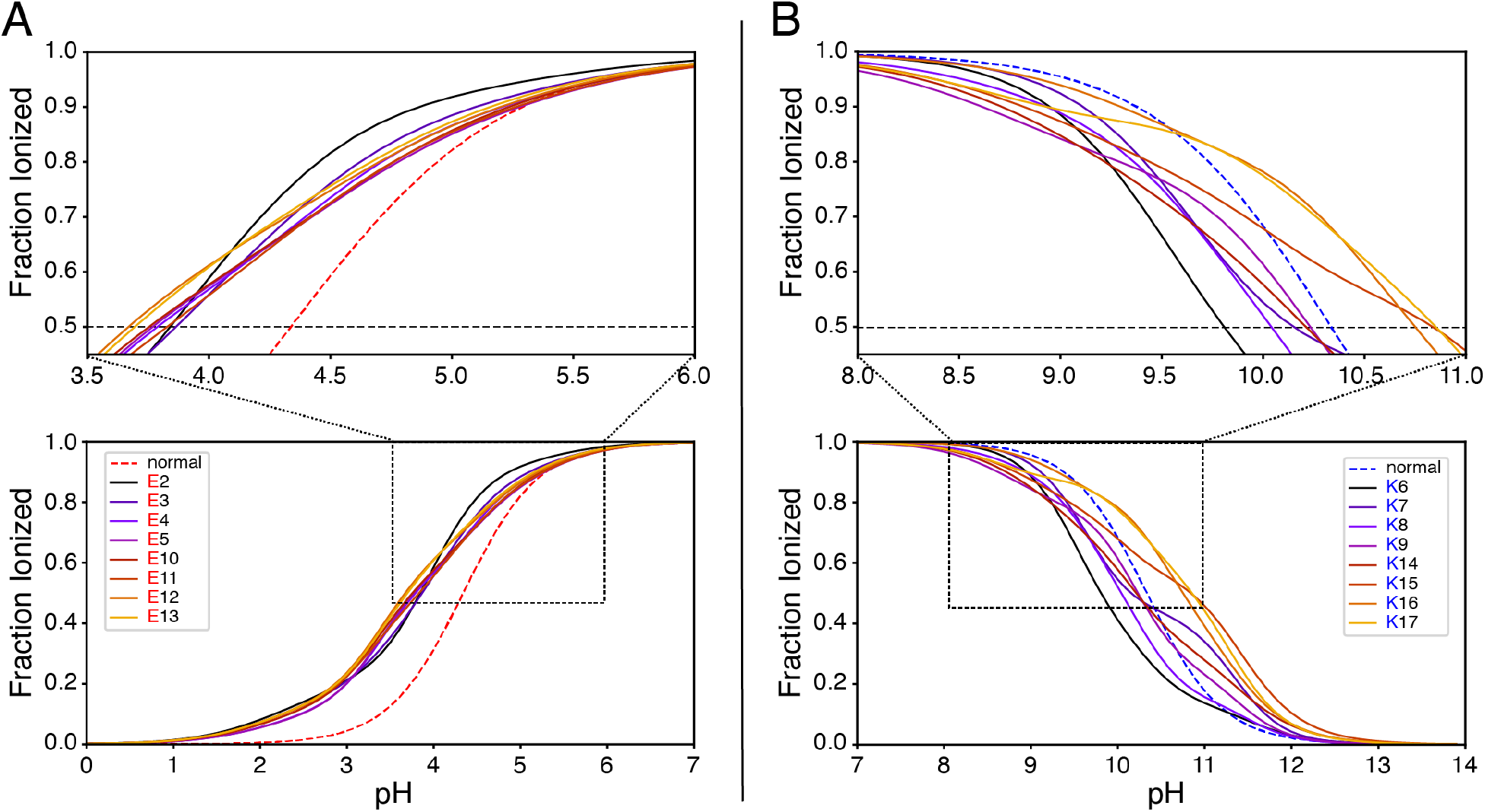
Site specific ionization curves for each of the positions in (E_4_K_4_)_2_. The curves are shown by dividing the pH range into two parts. This helps visualize the titration of Glu (panel A) and the titration of Lys residues (panel B). The top row shows zoomed in versions that focus on specific pH ranges to help with identifying the pH values at which the likelihood of being charged is 0.5 for the different residues (see black dashed lines). The red and blue dashed curves correspond to the ideal case of ionizable residues titrating at pK_a_ values corresponding to those of model compounds (10.34 for Lys and 4.34 for Glu ^52^), which in the legend is referred to as “normal”.

Conversely, the Lys residues show greater heterogeneity in terms of their apparent pK_a_ values. There can be down or upshifts of the apparent pK_a_ values when compared to the model compound value of 10.34 ^52^. The dispersion of apparent pK_a_ values is greater than one pH unit.

To quantify the preferential, site-specific protonation of ionizable residues, we introduce a parameter Ω_*q*,*r*_ for each site *r* along the sequence. Within a given mesostate of net charge *q*, the Ω_*q*,*r*_ parameter is defined as:

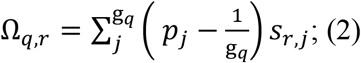

Here, Ω_*q*,*r*_ is the preferential protonation of residue *r* in a mesostate with net charge *q*; *p_j_* is the probability associated with microstate *j*; g_*q*_ is the number of microstates within the mesostate of net charge *q*, and *s_r,j_* is a switch function that quantifies the state of residue *r* in microstate *j*. The switch function has a value 1 if the residue is charged, and 0 if it is uncharged. The difference, shown in parentheses of the summand quantifies the difference between the microstate probability *p_j_*, derived from the properly reweighted ensemble and (*g_q_*)^−1^, which is the value we would expect from the postulate of equal *a priori* probabilities, the null model which sets all microstates within a mesostate to be equiprobable. If the summand is positive, it means that the microstate in question has a higher probability of being charged when compared to the null model. Conversely, if the summand is negative, then the microstate *j* has a lower probability of being charged when compared to the null model.

Using the information for all Ω_*q*,*r*_ values computed across all mesostates, we obtain the pH-dependent preferential protonation values per residue. This profile is denoted as Ω_*r*_(pH). It is computed as a weighted sum over mesostates populations at a given pH and is written as:

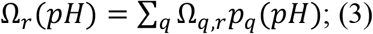

The results for Ω_*r*_(pH) at each position *r* are plotted in **Figure 9**. The results are shown for each of the three systems at pH 7.5. Data for each system are shown along the columns. Each row shows the Ω_*r*_(pH) profiles computed for the reweighted ensemble (top row), the helical ensemble (middle row), and the disordered ensemble (bottom row).

**Figure 9:**
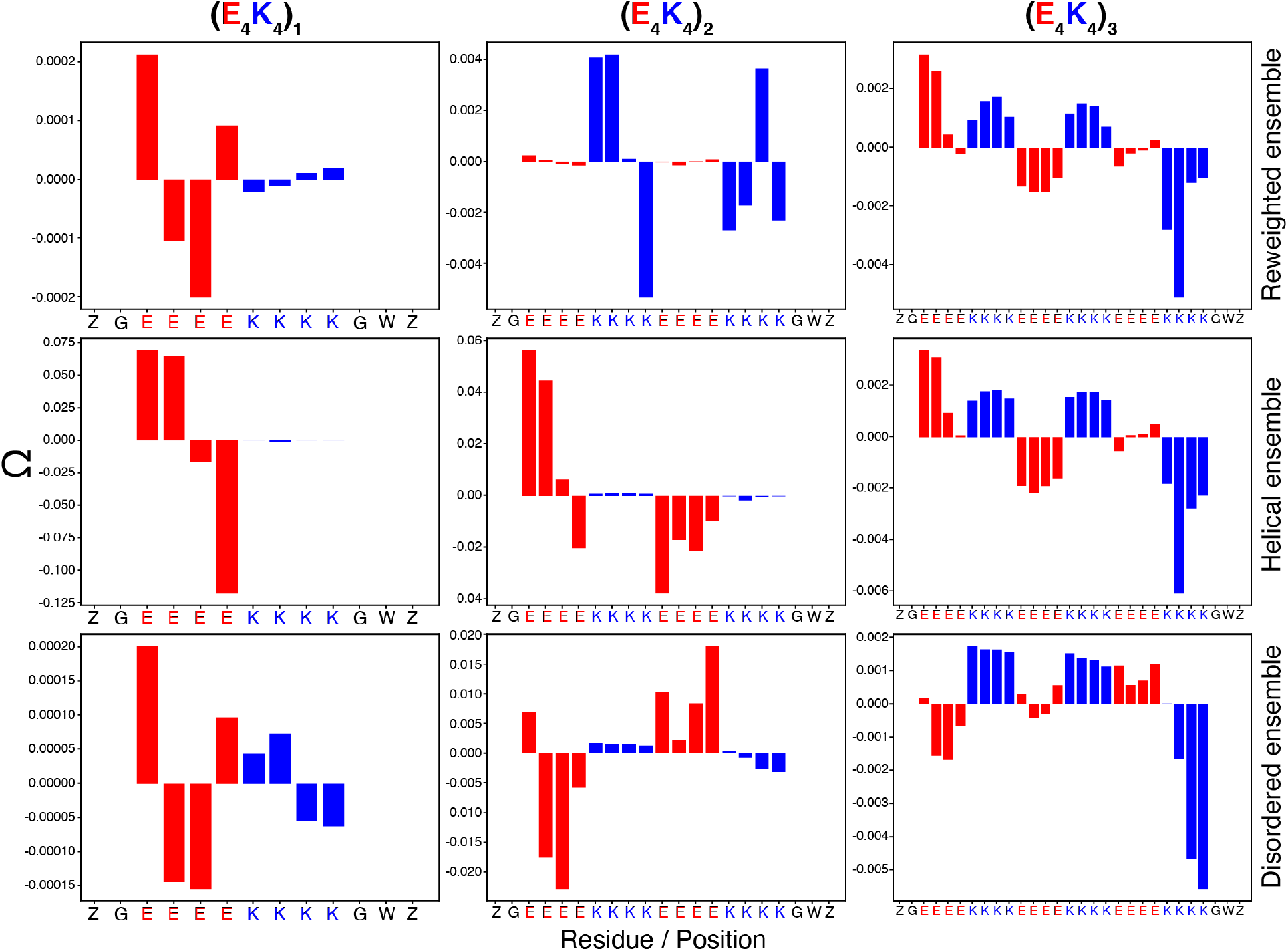
Ω profiles at pH 7.5 for each of the three peptides, organized along the columns, and types of ensembles, organized along the rows. The definition of Ω is in the main text. Positive values for Ω indicate a higher propensity for that residue to be in its charged form when compared to the scenario where all residues of the same type have the same pK_a_ value. Negative values of Ω indicate a higher propensity for that residue to be in its uncharged form when compared to the scenario where all residues of the sane type have the same pK_a_ value. Along the abscissa, the Z at the N-terminal end is the label for N-acetyl and Z at the C-terminal end is for NH2. We use Z as a single letter code for both caps. The Ω values increase significantly in magnitude for pH values that are above or below 7.5. We focus here on pH 7.5 to show the non-uniformity of preferred charge states at distinct positions. Further, Ω is calculated by referencing to a prior that accounts for the full degeneracy of microstates per mesostate. The magnitudes of Ω values become significantly larger if we were to use a prior based on fixed charge states whilst ignoring the degeneracy of microstates.

Positive values for Ω_*r*_(pH) imply that there is a preference to be charged, more so than would be expected based on the assumption of a fixed pK_a_ value for all residues of the same type. Conversely, negative values for Ω_*r*_(pH) imply that there is a preference to be uncharged, more so than would be expected based on the assumption of a fixed pK_a_ value for all residues of the same type. Note that these interpretations for the positive vs. negative deviations come from the sign of the summand in Equation (2). It depends on the microstate probability as inferred from the simulations. These probabilities are referenced to the probability we would compute by assuming that all microstates within a mesostate are equiprobable.

For the longest sequence, (E_4_K_4_)_3_ in the reweighted or fully helical ensembles, there is a higher preference for the internal Glu residues to be uncharged, when compared to the model that assumes fixed charged states. The converse is true for the internal Lys residues. In the reweighted and fully helical ensembles, there is an increased preference for charged Glu residues at the N-terminal capping position and an increased preference for uncharged Lys residues at the C-terminal capping position. The Ω profiles are inverted in the disordered ensemble (**Figure 9**).

Overall, the observations summarized in **Figure 9** can be traced to a series of factors that include the following: (i) The intrinsic helical propensity of Glu° is higher than that of Glu^− 28^. In fact, the intrinsic helical propensity of Glu° can even be higher than that of alanine, which is typically used as the standard for high intrinsic helical propensity ^28^. This higher intrinsic helical propensity helps rationalize the increased preference for Glu° at internal positions. Additionally, Ermolenko et al., showed that the helical propensities of uncharged residues are independent of their positions within the middle of alpha helices ^53^. This provides further support for our observation of preferential substitutions of Glu^−^ with Glu° in internal positions of sequences with (E_4_K_4_) repeats. (ii) Our finding of a clear positional preference for the charged Glu residues at the N-terminal end is concordant with documented preferences for Glu^−^ at the N-terminal caps, and the so-called N2 position ^12, 13^. Our observation of a positional preference for uncharged Lys° residues at the C-terminal caps is concordant with documented statistical preferences for polar amino acids and even hydrophobic groups at C-terminal capping positions ^12, 13, 54^. (iii) In the disordered ensembles, we observe a inversion, with one exception, of the positional preferences for being charged vs. uncharged. This reflects the dominance of local sequence context effects and contributions of conformational heterogeneity in the disordered ensembles. These effects engender a preference for maintaining Lys residues in their charged states at internal positions. Further, the increased preference of C-terminal Lys residues to be uncharged is also present in the disordered ensemble.

### There is a finite probability of observing uncharged residues instead of Glu or Lys residues in naturally occurring SAH sequences

Our findings suggest that maintenance of helical conformations across a wide range of pH values comes with a significant increase in charge state heterogeneity as the pH deviates from 7.5. This is likely to be become more pronounced in longer SAH sequences given the combinatorial increase in the number of charge states. One way to reduce this increased charge state heterogeneity would be to substitute internal Glu or Lys residues with non-ionizable residues. We quantified this probability across numerous sequences predicted to form single alpha helices (**Figure 10**). Overall, the likelihood of finding a non-ionizable residue at an internal position is only slightly lower than the likelihood of finding a Glu or Lys. This implies that there is an evolutionary drive to reduce some of the charge state heterogeneity that comes with the combinatorial increase of the numbers of charge states as the lengths of SAH forming sequences increase. We also observe a finite, albeit small likelihood that a Glu or Lys residue at an internal position will be occupied by a residue of the opposite charge. It is likely that these charge inverting interruptions disrupt the propagation of the SAH motif, again highlighting the need for caution in designating these systems as “single” alpha helix forming sequences.

**Figure 10:**
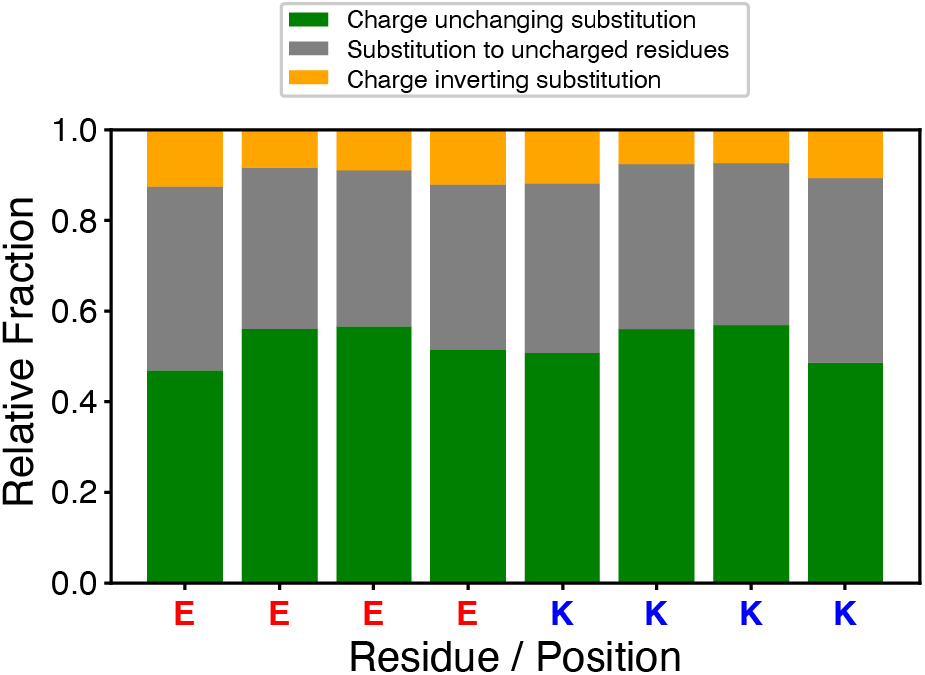
Stacked bar plots showing the substitution patterns for Glu and Lys residues within internal repeats across 240,754 predicted SAH sequences extracted from the CSAH database ^55^. Conservation implies that the charge at a specific position within the idealized repeat is preserved when compared to the consensus repeat. Substitution to uncharged residues quantifies the fraction of residues in that position in the idealized repeat that have been substituted with uncharged residues in naturally occurring sequences. Finally, charge inversion quantifies the observed frequency of substituting residues in each position of the idealized repeat whereby the substitution inverts the charge. The database contains all predictions on the Trembl part of the uniprot database ^56^.

## DISCUSSION

Motivated by the pH insensitivity of measured helicity in SAH forming sequences ^34^ (**Figure 2**), we have used our formulation of the *q*-canonical ensemble ^29^ to investigate the effects of charge state heterogeneity and charge regulation. Our studies were made possible by decoupling charge and conformational measurements, and using the information derived from the two sets of independent measurements as restraints for reweighting atomistic simulations.

The picture that emerges is as follows: There are several microstates and hence mesostates with a net charge that have intrinsic helicities that are equivalent to or higher than those of the single microstate within the 0_1_ mesostate that has a net charge of zero. The identification of mesostates with high intrinsic helicities comes from joint analysis of the potentiometric titrations and UV-CD measurements as a function of pH. The relevant microstates within each mesostate are obtained from atomistic simulations that use the ABSINTH implicit solvation model and *q*MC sampling ^50^.

There is a clear increase in the number of microstates and hence mesostates that contribute to the *q*-canonical ensemble as the number of repeats increase. Overall, the ensembles display high charge state heterogeneity and low conformational heterogeneity, with helical conformations dominating the ensemble. Even at pH 7.5, there is a finite, albeit small likelihood that the (E_4_K_4_)_3_ system can sample microstates with a net charge of ±1. As shown in **Figure 4**, the intrinsic helicities of these mesostates with a net charge are equivalent to those of the 0_1_ mesostate. For the SAH forming systems, several charge microstates / mesostates are compatible with forming stable helical conformations. This charge state heterogeneity or diversity buffers against large-scale conformational changes – a phenomenon we refer to as *conformational buffering*. The term itself is coopted from the recent work of González-Foutel et al.,^57^.

Why might conformational buffering combined with charge state heterogeneity be important in biology? Because a given SAH sequence can adopt multiple charge states whilst sampling very similar conformational ensembles. This suggests that the interactions mediated by an SAH sequence will be charge state dependent, and hence diverse, even though the conformational template remains essentially the same. Accordingly, the effects of charge regulation can engender a sort of *many from one* form of molecular recognition, whereby a single SAH sequence can enable the recognition of diverse substrates of different charge profiles.

We find that preferential protonation profiles at a given pH (**Figure 9**) are markedly different for the helical vs. disordered ensembles. This highlights how position-specific proton binding and release combined with the charge state heterogeneity can tilt conformational biases toward specific types of conformations. In effect, one can think of helix formation in so-called SAH sequences as a simultaneous minimization of enthalpy due to helix formation ^17, 21^ and maximization of sequence entropy through charge state heterogeneity. Our analysis of frequencies of replacement of Glu or Lys residues with uncharged residues (**Figure 10**) suggests that some of the charge state heterogeneity may be reduced for specific sequences in the SAH superfamily, and this reduction might be necessary for the different functional contexts in which specific SAH-forming elements are used.

From a physico-chemical perspective, we propose that there are three main factors that contribute to the linkage between proton binding and conformational equilibria appear to be as follows: (i) The intrinsic conformational biases of the neutral vs. charged forms of ionizable residues ^28, 58^. (ii) The intrinsic ^41^ and sequence / conformational context dependence of the free energies of solvation of neutral vs. charged forms of ionizable residues. And (iii), the interplay between local and non-local electrostatic attractions vs. repulsions ^59^. In isolated alpha helical conformations such as the SAH, the functional groups of ionizable residues can be pushed out from the backbone, thus maximizing the extent of solvation ^22^. This is important because the free energies of solvation of charged forms of ionizable residues are at least an order of magnitude more favorable than the uncharged forms ^41, 46^. However, within helical conformations, there is likely to be an overlap of solvation shells of the Lys^+^ and Glu^−^ residues ^23, 60^. These overlaps are likely to be thermodynamically unfavorable because the water molecules within the overlapping solvation shells need to be oriented differently around amines vs. carboxyl groups ^41^. Selective, position-specific proton binding or release can alleviate orientational restrictions by minimizing the overlap of solvation shells. (iii) The high local charge density engenders the possibility of local repulsions. These will need to be minimized while enabling favorable *i* to *i*+4 salt bridges to form.

Overall, our findings suggest that some level of charge regulation, the extent of which depends on pH, will contribute to the complex determinants of SAH stability. This observation is consistent with the realization that the determinants of helix stability can be quite complex ^61^. There is always an intricate interplay between the enthalpy and entropy of the peptide plus solvent system, and how this is achieved can be highly context specific.

SAHs are typically viewed as rod-like semi-flexible entities ^37^. This has led to the deployment of SAH sequences as rigid molecular rulers for various intracellular conformational measurements ^39^. In these measurements, the number of repeats is used as a direct route to change the scale of the ruler. This would be optimal if and only if the assumption of a rigid, rod-like conformation were valid, and that each of the (E_4_K_4_)_n_ peptides were 100% alpha helical. However, this is not true even for a two-state equilibrium where one assumes a perfect, rod-like conformation in equilibrium with a coil-like ensemble. Further, the assumptions of rigid, rod-like conformations become more concerning when one realizes that helical rod is unlikely to be the dominant conformation for peptides such as the (E_4_K_4_)_3_ systems studied here (**Figure 7**). Of course, one might be inclined to critique the observed conformational heterogeneity we report in Figure 7. Indeed, recent work suggests that the failure to observe a rod-like alpha helical conformation in 100% of the ensemble represents a failure of a specific brand of implicit solvation models ^62^. In that study, the authors set up the expectation that (E_4_K_4_)_n_ systems should be nearly 100% alpha helical and perfectly rod-like. This goes against what is known and well established in the helix-coil transition literature. The helical rod requires that the sequence length be infinitely long. In practical terms, the length must be closer to or beyond the higher end of the observed length distribution of most SAH forming sequences ^35^ (**Figure S1**). Simulations based on explicit representations of solvent molecules do not show reversible transitions between helical and non-helical conformations ^33^. Instead, they have been used for understanding the role of hydration in mediating *i* to *i*+4 salt bridges. Therefore, simulations based on classical forcefields, irrespective of whether they are based on implicit vs. explicit solvation models, are likely to be confounded by site- and context-specific protonation effects. These may be accountable in next generation polarizable models such as the HIPPO model that accounts for polarizability i.e., electron cloud deformation, and the effects of charge transfer ^63^. Whether such models can be made efficient and deployed to account for charge regulation effects is unclear at this juncture. For now, the practical approach of combining information from experiments that decouple measurements of charge and conformation with simulations that explicitly account for charge state heterogeneity, best achieved using a well parameterized implicit solvation model such as ABSINTH, is, at least from our perspective, the optimal route to pursue for uncovering the effects of charge regulation. The ABSINTH framework includes the explicit representation of solution ions, and this will be useful for dissecting the interplay between charge regulation and charge renormalization (see below).

It is well-known that SAH forming sequences can be misclassified as being intrinsically disordered or as drivers of coiled coils ^31, 38^. This raises the formal possibility that some fraction of sequences that are predicted to be intrinsically disordered are likely to form well-ordered conformations. In such systems, especially those enriched in charged residues, there might indeed be low conformational heterogeneity that serves as a buffer against high charge state heterogeneity. This could also be true of regions predicted to be disordered that are also substrates of multisite post-translational modifications such as Ser / Thr phosphorylation, Lys acetylation, Arg methylation or citrullination ^64^. What we need are combined interrogations of charge states using potentiometry, conformational states using multipronged spectroscopies, and simulations that account for heterogeneous distributions of charge states and the conversions between these states^50^. We have presented one such method here and summarized the insights that emerge from this combined methodology.

Finally, in addition to charge regulation effects, there are likely to be *charge renormalization* effects due to preferential territorial organization of solution ions around ionizable groups ^65, 66^. These effects are likely to be minor at least for the 0_1_ mesostate, although a detailed investigation of the interplay of charge regulation ^29^ and charge renormalization ^67^ requires deeper scrutiny given that the cellular milieu is a complex mixture of protons, ions, and other solutes ^65, 68^.

## MATERIALS AND METHODS

### Materials

Potassium chloride (KCl), hydrochloric acid (HCl), potassium hydroxide (KOH), and potassium hydrogen phthalate (KHP) were purchased from Sigma (St. Louis, MO, USA). We assessed the purity of KOH using titrations of a standard with a known pK_a_, which confirm that contaminants, if present are miniscule and do not affect the potentiometric titrations^29^. Peptides were purchased from GenScript (Piscataway, NJ, USA) at >95% purity as HCl salts with acetylated N-termini and amidated C-termini to avoid confounding effects from charged termini. Salt content was determined by mass spectrometry. Peptides were stored in lyophilized form at −20°C in sealed containers in the presence of desiccant until used for experiments. Three peptides were used in this study: ac-G-(E_4_K_4_)_n_-GW-NH_2_, where n = 1, 2, or 3. Tryptophan was used to enable precise measurements of concentration.

### Potentiometry

Potentiometric titrations were carried out in CO_2_-depleted solutions as previously described in an earlier report ^29^. The data used here are identical to those in the previously published work ^29^.

### UV-CD measurements

Samples were prepared for UV-CD measurements by carrying out a potentiometric titration as previously described ^29^, except that at regular increments throughout the titration, 40 μL aliquots were removed from the sample being titrated and set aside for UV-CD measurements. Subsequently, these aliquots were loaded sequentially into a 0.1 mm pathlength demountable quartz cuvette, and UV-CD spectra were collected at 20°C using a JASCO J-810 spectropolarimeter. All UV-CD spectra were the average of six scans from 260-190 nm, with a 1 nm step, a 2 second response time, and 50 nm/min scanning speed. The UV-CD spectra measured for background solution at low and high pH were identical within error, so the low pH background was subtracted from all spectra. The resulting ellipticity was used to calculate the mean residue ellipticity ([θ], units of deg cm^2^/dmol residue) using Equation (3):

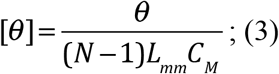

where θ = ellipticity (mdeg, machine units), *N* is the number of residues, *N*–1 is the number of backbone amides, *L*_mm_ is the path length of the cuvette (mm), and *C_M_* is the protein concentration in molar units.

### Deconvolution of UV-CD data

Data were converted to units of mean residue ellipticity as described above and then analyzed by deconvolution to obtain estimates of secondary structure content. Deconvolution analysis was carried out using the CDSSTR algorithm as implemented through the Dichroweb server. Basis sets 4, 7, and SP175 were used for the deconvolution, as these are the basis sets that have been optimized for the wavelength rage of 190-240 nm. The CDSSTR algorithm consistently provided the best fits for our data and outperformed other deconvolution algorithms including CONTIN and SELCON3. The outputs of the deconvolution analysis are reported as fractional secondary structure content in six categories: alpha helix regular, alpha helix distorted, beta sheet regular, beta sheet distorted, turns, and unordered. For the sake of simplicity, we sum the two helical categories to obtain a single estimate of fractional helicity. The secondary content estimates reported in this work represent the mean and standard deviation of the values resulting from analysis using each of the three basis sets (4, 7, SP175).

### Extracting intrinsic helicities of mesostates

We recently published a protocol to extract the probabilities of individual mesostates as a function of pH ^29^. As noted in Equation (1), the overall helicity profile as a function of pH is the weighted sum of the convolution of the probability of accessing a mesostate of net charge *q* at a given pH and the intrinsic helicity associated with the mesostate. The latter, as noted in Equation (1), is independent of pH and intrinsic to the mesostate. The intrinsic helicities of mesostates are inferred through a Monte Carlo based numerical fitting procedure designed to reproduce the measured helicity profile, which is denoted by the left-hand side of Equation (1). The mesostate populations as a function of pH are known from potentiometric titrations. Accordingly, we define a cost function as the difference in between the simulated curve and the experimental profile:

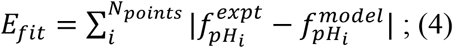

Here, *pH_i_* is the pH at which the data point of interest was collected, 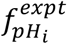 is the experimental data point, and 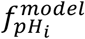 is the value for the fractional helicity that we estimate at *pH_i_* from the current set of intrinsic helicities of mesostates. The latter are chosen by sampling at random, in an interval between 0 and 1 and accepting or rejecting the proposed move in parameter space using the Metropolis-Hastings criterion defined in Equation (5) as:

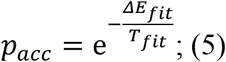

Here, *p_acc_* is the acceptance probability of a specific move to increase or decrease the fraction helicity, *ΔE_fit_* is the difference in fitting energy between the last accepted move, and the current proposed move. This step is repeated for 10^4^ steps and each Monte Carlo sweep involving 10^4^ steps was performed separately and restarted 10^4^ times.

### ABSINTH Simulations

All the simulation were performed using the ABSINTH implicit solvent model ^49, 69^. We used the *q*-canonical Monte Carlo sampling method ^50^, with a few modifications. Instead of using the populations of each microstate within a single mesostate simulation to obtain the standard free energy differences, we performed individual simulations for each of the microstates. The temperature for these simulations as set to be 240 K for the unmodified ensemble, 350 K for the disordered ensemble, and 278 K for the helical ensemble. These simulation temperatures are the same for all repeat lengths. To generate the final ensemble, we set all temperatures to the experimental temperature of 298 K and mix conformations drawn from the three different ensembles. For each microstate, the ABSINTH-based Monte Carlo simulations use at 10^8^ steps. Information from the first set of 5×10^6^ steps was not used in the analysis, as this was the equilibration phase in the simulations. The conformation and frame specific energies were saved every 10^4^ steps. We then derive the free energy differences between microstates by performing cross energy calculations. These calculations entail reassessing the energy for each frame obtained for a given microstate while substituting the potential of a different microstate. The cross energies were then used as input for the Bennett Acceptance Ratio (BAR) procedure, as implemented in the python package pymbar ^70^.

### Reweighting of the simulations

We use the helicity per mesostates and mesostate populations as a function of pH to reweight the overall ensemble by appropriate mixing of the three simulated ensembles. The three ensembles are then mixed, and mixing is the reweighting procedure that is designed to reproduce the intrinsic helicities of individual mesostates. For each mesostate, we start by assessing the overall helicity in each of the three sets. We then compare the mesostate helicities of the unmodified set to the helicity per mesostate that we obtained from experiments. If the helicity is too high for that mesostate compared to the experimentally derived value, we introduce the first set until the overall weighted sum of the two sets are within 0.001 of experiments. Conversely, if the helicity is too low for that mesostate compared to the experimentally derived value, we introduce the third set to find the overall ensemble. The proportion of mixing of each ensemble is shown in **Figure S6**. Using the mixing function, we combined all properties of each microstate within each mesostate to match experiments. We then set the mesostate free energies to the free energies obtained from potentiometric data, while conserving the properly mixed microstate weights within mesostates. This means that the average free energy of each mesostate is the potentiometrically derived values, but the free energy differences within mesostates are those derived from simulations, after proper mixing. This allows us to extract microstate weights that are consistent with experiments, while also generating structural insights on a per residue basis. Because of the large number of simulations needed to get the full pH dependence in (E_4_K_4_)_3_, we focused on the mesostates with net charge ranging from −3 to 3. This is reasonable because potentiometry shows that these mesostates contribute to more than 99% of overall population across pH values between 5.3 to 9.7 ^29^. For the constructs of (E_4_K_4_)_1_ and (E_4_K_4_)_2_ repeats, the computational cost is significantly lower, and we were able to obtain simulations for every microstate. The simulations were analyzed using the MDtraj python package ^51^. This includes determination of the secondary structure contents, quantification of the numbers of salt bridges, etc. In our analysis, a salt bridge was approximated using a simple average of the number of counts of opposite charge ionized residue pairs within 4 Å of each other.

## ASSOCIATED CONTENT

The supplemental material includes Figures S1 – S8.

## AUTHOR INFORMATION

### Author Contributions

All authors designed the study. M.J.F., developed the *q*-canonical ensemble framework. M.J.F., designed the simulation setup, performed the simulations, and analyzed the simulation results. A.E.P., designed and deployed the potentiometric titrations and performed all the measurements. M.J.F., A.E.P., and R.V.P., analyzed all the data. All authors contributed to the writing and editing of the manuscript. R.V.P., secured funding.

## ACKNOWLEDGMENTS

This work was supported by grants from the Air Force Office of Scientific Research (FA9550-20-1-0241), the US National Institutes of Health (R01NS121114), and the St. Jude Children’s Research Hospital. We thank Mina Farag, Matthew King, Min Kyung Shinn, and Xiangze Zeng for helpful discussions and comments on the manuscript.

## REFERENCES

[1] L. Pauling, R. B. Corey, H. R. Branson Proceedings of the National Academy of Sciences. 1951, 37, 205–211.

[2] D. Eisenberg Proceedings of the National Academy of Sciences. 2003, 100, 11207–11210.

[3] W. Kabsch, C. Sander Biopolymers. 1983, 22, 2577–2637.

[4] B. H. Zimm, J. K. Bragg The Journal of Chemical Physics. 1959, 31, 526–535.

[5] S. Lifson, A. Roig The Journal of Chemical Physics. 1961, 34, 1963–1974.

[6] D. C. Poland, H. A. Scheraga The Journal of Chemical Physics. 1965, 43, 2071–2074.

[7] A. Vitalis, A. Caflisch Journal of chemical theory and computation. 2012, 8, 363–373.

[8] R. Aurora, T. P. Creamer, R. Srinivasan, G. D. Rose Journal of Biological Chemistry. 1997, 272, 1413–1416.

[9] S. Marqusee, V. H. Robbins, R. L. Baldwin Proceedings of the National Academy of Sciences. 1989, 86, 5286–5290.

[10] S. Padmanabhan, S. Marqusee, T. Ridgeway, T. M. Laue, R. L. Baldwin Nature. 1990, 344, 268–270.

[11] A. Chakrabartty, R. L. Baldwin in Stability of α-Helices, Vol. 46 (Eds.: C. B. Anfinsen, F. M. Richards, J. T. Edsall, D. S. Eisenberg), Academic Press, 1995, pp.141–176.

[12] A. J. Doig, R. L. Baldwin Protein science : a publication of the Protein Society. 1995, 4, 1325–1336.

[13] R. Aurora, G. D. Rosee Protein science : a publication of the Protein Society. 1998, 7, 21–38.

[14] P. Robustelli, S. Piana, D. E. Shaw Journal of the American Chemical Society. 2020, 142, 11092–11101.

[15] K. Shiraki, K. Nishikawa, Y. Goto Journal of molecular biology. 1995, 245, 180–194.

[16] J. K. Myers, C. Nick Pace, J. Martin Scholtz Protein science : a publication of the Protein Society. 1998, 7, 383–388.

[17] M. M. Lopez, D.-H. Chin, R. L. Baldwin, G. I. Makhatadze Proceedings of the National Academy of Sciences. 2002, 99, 1298–1302.

[18] K. Ghosh, K. A. Dill Journal of the American Chemical Society. 2009, 131, 2306–2312.

[19] J. M. Rogers, C. T. Wong, J. Clarke Journal of the American Chemical Society. 2014, 136, 5197–5200.

[20] C. J. Oldfield, Y. Cheng, M. S. Cortese, P. Romero, V. N. Uversky, A. K. Dunker Biochemistry. 2005, 44, 12454–12470.

[21] S. Marqusee, R. L. Baldwin Proceedings of the National Academy of Sciences. 1987, 84, 8898–8902.

[22] H. Lu, J. Wang, Y. Bai, J. W. Lang, S. Liu, Y. Lin, J. Cheng Nature communications. 2011, 2, 206.

[23] M. Sundaralingam, Y. C. Sekharudu, N. Yathindra, V. Ravichandran Proteins: Structure, Function, and Bioinformatics. 1987, 2, 64–71.

[24] C. B. Stanley, H. H. Strey Biophysical Journal. 2008, 94, 4427–4434.

[25] D. S. Olander, A. Holtzer Journal of the American Chemical Society. 1968, 90, 4549–4560.

[26] M. Nagasawa, A. Holtzer Journal of the American Chemical Society. 1964, 86, 538–543.

[27] E. A. Gooding, S. Sharma, S. A. Petty, E. A. Fouts, C. J. Palmer, B. E. Nolan, M. Volk Chemical Physics. 2013, 422, 115–123.

[28] C. Nick Pace, J. Martin Scholtz Biophysical Journal. 1998, 75, 422–427.

[29] M. J. Fossat, A. E. Posey, R. V. Pappu Biophysical Journal. 2021, 120, 5438–5453.

[30] A. L. Boyle in 3 – Applications of de novo designed peptides, Vol. (Ed. S. Koutsopoulos), Woodhead Publishing, 2018, pp.51–86.

[31] D. Suveges, Z. Gaspari, G. Toth, L. Nyitray Proteins: Structure, Function, Bioinformatics. 2009, 74, 905–916.

[32] S. Iwata, J. W. Lee, K. Okada, J. K. Lee, M. Iwata, B. Rasmussen, T. A. Link, S. Ramaswamy, B. K. Jap Science (New York, N.Y.). 1998, 281, 64–71.

[33] M. Wolny, M. Batchelor, G. J. Bartlett, E. G. Baker, M. Kurzawa, P. J. Knight, L. Dougan, D. N. Woolfson, E. Paci, M. Peckham Scientific Reports. 2017, 7, 44341.

[34] E. G. Baker, G. J. Bartlett, M. P. Crump, R. B. Sessions, N. Linden, C. F. J. Faul, D. N. Woolfson Nature Chemical Biology. 2015, 11, 221–228.

[35] C. A. Barnes, Y. Shen, J. Ying, Y. Takagi, D. A. Torchia, J. R. Sellers, A. Bax Journal of the American Chemical Society. 2019, 141, 9004–9017.

[36] S. Iwai, D. Hanamoto, S. Chaen Journal of Biological Chemistry. 2006, 281, 30736–30744.

[37] J. A. Spudich, S. Sivaramakrishnan Nature Reviews Molecular Cell Biology. 2010, 11, 128–137.

[38] M. Peckham, P. J. Knight Soft Matter. 2009, 5, 2493–2503.

[39] R. U. Malik, M. Dysthe, M. Ritt, R. K. Sunahara, S. Sivaramakrishnan Scientific Reports. 2017, 7, 7749.

[40] E. Wang, C.-L. A. Wang Archives of Biochemistry and Biophysics. 1996, 329, 156–162.

[41] M. J. Fossat, X. Zeng, R. V. Pappu The Journal of Physical Chemistry B. 2021, 125, 4148–4161.

[42] C. A. Fitch, D. A. Karp, K. K. Lee, W. E. Stites, E. E. Lattman, E. B. Garcia-Moreno Biophysical Journal. 2002, 82, 3289–3304.

[43] C. D. Andrew, J. Warwicker, G. R. Jones, A. J. Doig Biochemistry. 2002, 41, 1897–1905.

[44] N. Errington, A. J. Doig Biochemistry. 2005, 44, 7553–7558.

[45] D. Simm, K. Hatje, M. Kollmar PloS one. 2017, 12, e0174639.

[46] X. Zeng, C. Liu, M. J. Fossat, P. Ren, A. Chilkoti, R. V. Pappu APL Materials. 2021, 9, 021119.

[47] N. Fili, C. P. Toseland International Journal of Molecular Sciences. 2020, 21, 67.

[48] J. Li, Q. Lu, M. Zhang Traffic. 2016, 17, 822–838.

[49] A. Vitalis, R. V. Pappu Journal of computational chemistry. 2009, 30, 673–699.

[50] M. J. Fossat, R. V. Pappu The Journal of Physical Chemistry B. 2019, 123, 6952–6967.

[51] Robert T. McGibbon, Kyle A. Beauchamp, Matthew P. Harrigan, C. Klein, Jason M. Swails, Carlos X. Hernández, Christian R. Schwantes, L.-P. Wang, Thomas J. Lane, Vijay S. Pande Biophysical Journal. 2015, 109, 1528–1532.

[52] G. Platzer, M. Okon, L. P. McIntosh Journal of Biomolecular NMR. 2014, 60, 109–129.

[53] D. N. Ermolenko, J. M. Richardson, G. I. Makhatadze Protein science : a publication of the Protein Society. 2003, 12, 1169–1176.

[54] D. N. Ermolenko, S. T. Thomas, R. Aurora, A. M. Gronenborn, G. I. Makhatadze Journal of molecular biology. 2002, 322, 123–135.

[55] Á. Kovács, D. Dudola, L. Nyitray, G. Tóth, Z. Nagy, Z. Gáspári Journal of Structural Biology. 2018, 204, 109–116.

[56] Y. Wang, Q. Wang, H. Huang, W. Huang, Y. Chen, P. B. McGarvey, C. H. Wu, C. N. Arighi, C. on behalf of the UniProt PLOS Biology. 2021, 19, e3001464.

[57] N. S. González-Foutel, J. Glavina, W. M. Borcherds, M. Safranchik, S. Barrera-Vilarmau, A. Sagar, A. Estaña, A. Barozet, N. A. Garrone, G. Fernandez-Ballester, C. Blanes-Mira, I. E. Sánchez, G. de Prat-Gay, J. Cortés, P. Bernadó, R. V. Pappu, A. S. Holehouse, G. W. Daughdrill, L. B. Chemes Nature Structural & Molecular Biology. 2022, 29, 781–790.

[58] C. N. Pace, G. R. Grimsley, J. M. Scholtz Journal of Biological Chemistry. 2009, 284, 13285–13289.

[59] R. K. Das, R. V. Pappu Proceedings of the National Academy of Sciences of the United States of America. 2013, 110, 13392–13397.

[60] M. Sundaralingam, Y. C. Sekharudu Science (New York, N.Y.). 1989, 244, 1333–1337.

[61] J. M. Richardson, M. M. Lopez, G. I. Makhatadze Proceedings of the National Academy of Sciences. 2005, 102, 1413–1418.

[62] E. J. M. Lang, E. G. Baker, D. N. Woolfson, A. J. Mulholland Journal of chemical theory and computation. 2022, 18, 4070–4076.

[63] J. A. Rackers, R. R. Silva, Z. Wang, J. W. Ponder Journal of chemical theory and computation. 2021, 17, 7056–7084.

[64] G. Harauz, A. A. Musse Neurochemical Research. 2007, 32, 137–158.

[65] M. Krishnan The Journal of Chemical Physics. 2017, 146, 205101.

[66] R. R. Netz, H. Orland The European Physical Journal E. 2003, 11, 301–311.

[67] G. Kloes, T. J. D. Bennett, A. Chapet-Batlle, A. Behjatian, A. J. Turberfield, M. Krishnan Nano Letters. 2022, 22, 7834–7840.

[68] R. Milo, P. Jorgensen, U. Moran, G. Weber, M. Springer Nucleic Acids Research. 2009, 38, D750–D753.

[69] A. Mittal, R. Das, A. Vitalis, R. V. Pappu in ABSINTH Implicit Solvation Model and Force Field Paradigm for Use in Simulations of Intrinsically Disordered Proteins., Vol. (Ed. M. Fuxreiter), CRC Press (Taylor and Francis Group), Boca Raton, RL, 2015, pp.181–203.

[70] M. R. Shirts, J. D. Chodera Journal of Chemical Physics. 2008, 129, 124105.

