## Supplemental Material for "Uncovering the contributions of charge regulation to the stability of single alpha helices"

[†]Equal contributors

### Supplementary Figures

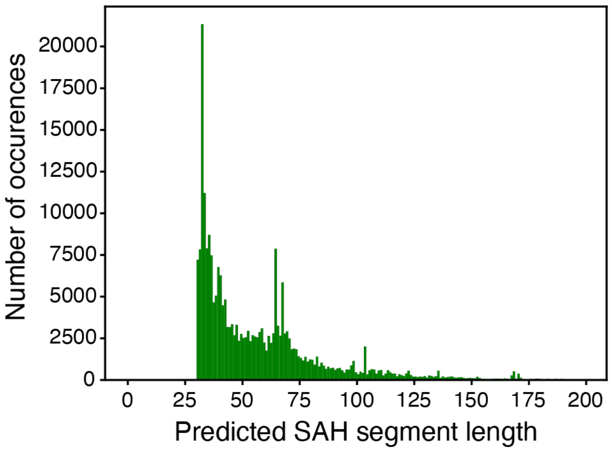


**Figure S1. Distribution of predicted Single Alpha Helices segment length from the 240754 Trembl entries of the CSAH database[1] predicted by the consensus of two algorithms**. SCAN4CSAH and FT_CHARGE[2,3]. We used the CSAH database http://csahserver.itk.ppke.hu/csah/download.html.

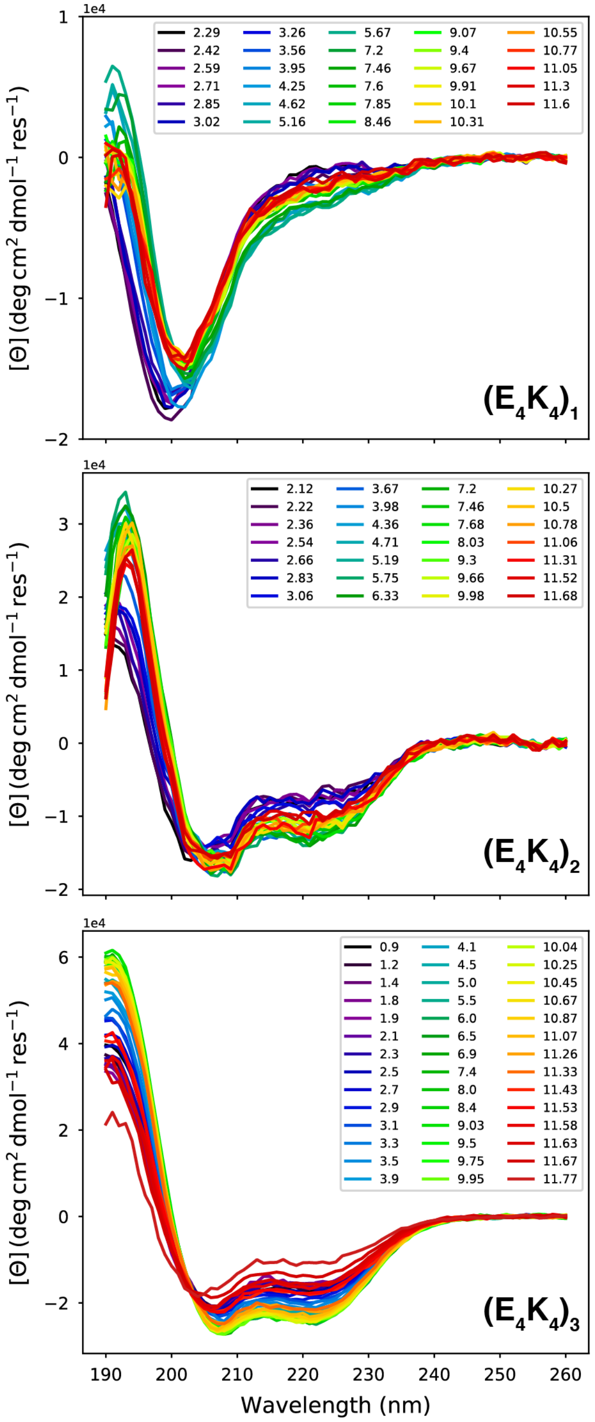


**Figure S2. CD spectra as a function of pH for ($E_4K_4$)$_1$ (top), ($E_4K_4$)$_2$ (middle), and ($E_4K_4$)$_3$ (bottom).** Samples were titrated as described for potentiometry experiments. CD spectra were collected in a 0.1 mm demountable quartz cuvette at 20°C.

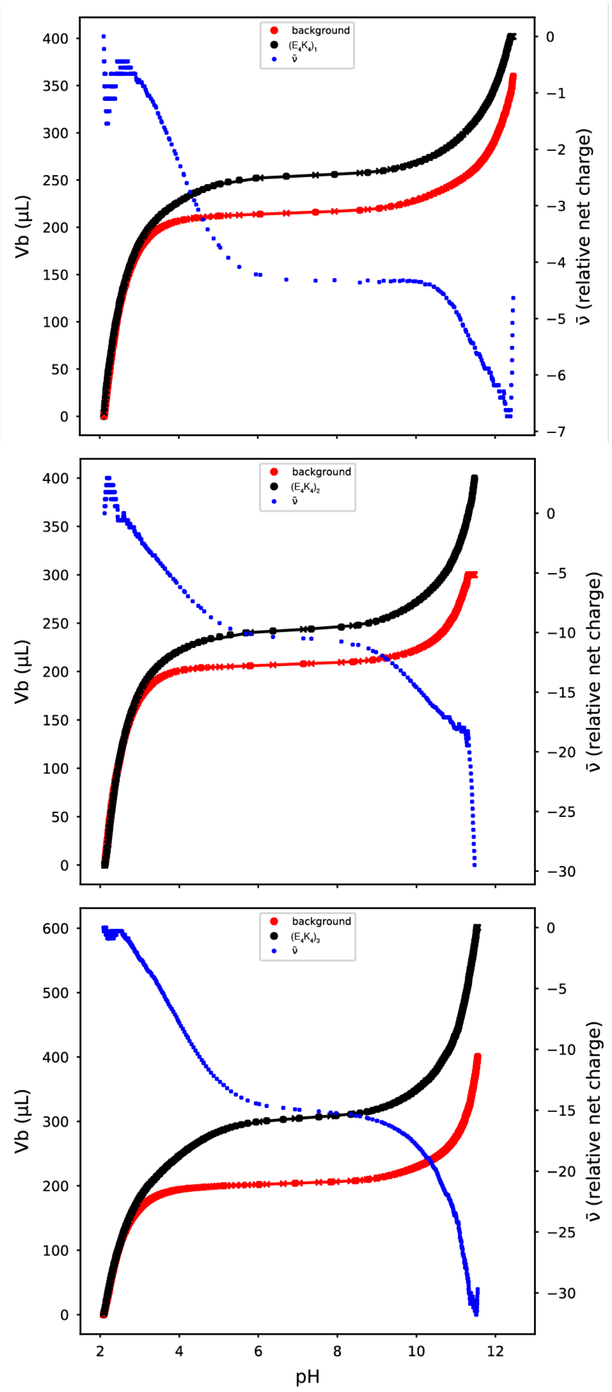


**Figure S3. Potentiometric measurements for $(E_4K_4)_1$ (top), $(E_4K_4)_2$ (middle), and $(E_4K_4)_3$ (bottom).** Titration data were transposed using the approach of Nozaki and Tanford to facilitate comparison of the volume of KOH added in the background titration (red circles) vs. the peptide titration (black circles) at each pH value. Additional points (-x-) were calculated via linear interpolation between data points to facilitate subtraction of the two curves at each pH value. The moles of protons released per moles of peptide, which is equivalent to relative change in net charge (blue points), were calculated from the difference of the red and black data points. The concentration of $(E_4K_4)_1$ was 215 μM and the titrant (KOH) concentration was 48.3 mM. The concentration of $(E_4K_4)_2$ was 95 μM and the titrant (KOH) concentration was 56.0 mM. The concentration of $(E_4K_4)_3$ was 180 μM and the titrant (KOH) concentration was 53.4 mM. This figure was adapted from Fossat et al. SI [4]

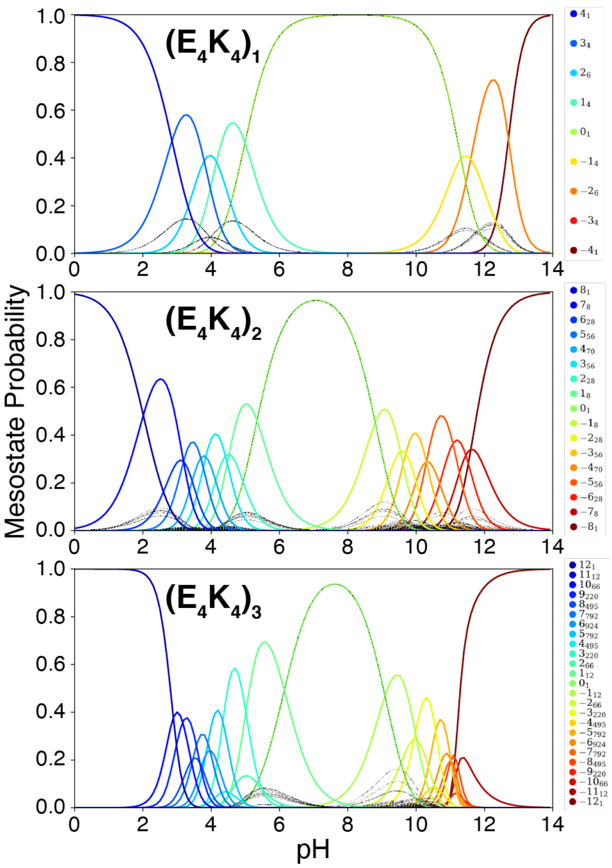


**Figure S4**: **Mesostate probabilities, plotted against pH, for the $(E_4K_4)_1$, $(E_4K_4)_2$, and $(E_4K_4)_3$ peptide. The colored lines correspond to the different mesostates, from most positive (blue) to most negative (red).** The dashed black line corresponds to the microstate probabilities constituting a mesostates, as obtained from the simulations. This figure was adapted from Fossat et al.[4].

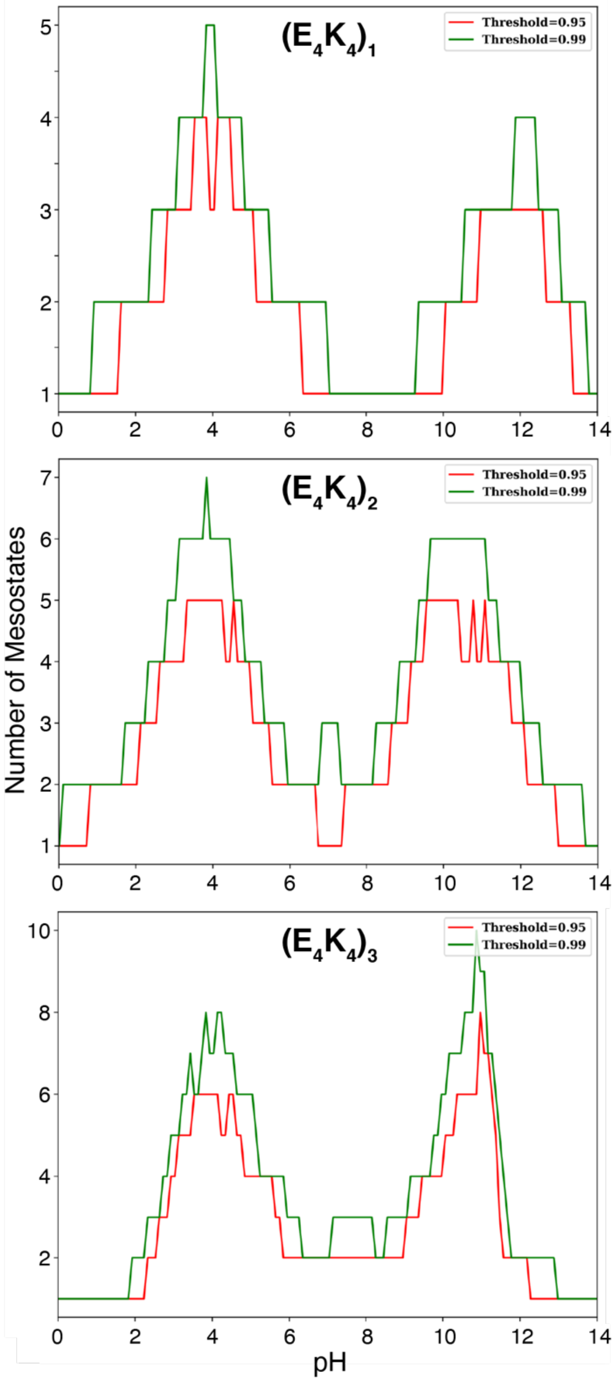


**Figure S5 : Minimum number of mesostates that are required to account for 95 and 99% of the population, as a function of pH, for each of the three peptides: $(E_4K_4)_1$ (A), $(E_4K_4)_2$ (B), and $(E_4K_4)_3$ (C).** These figures are from the work of Fossat et al.,[4].

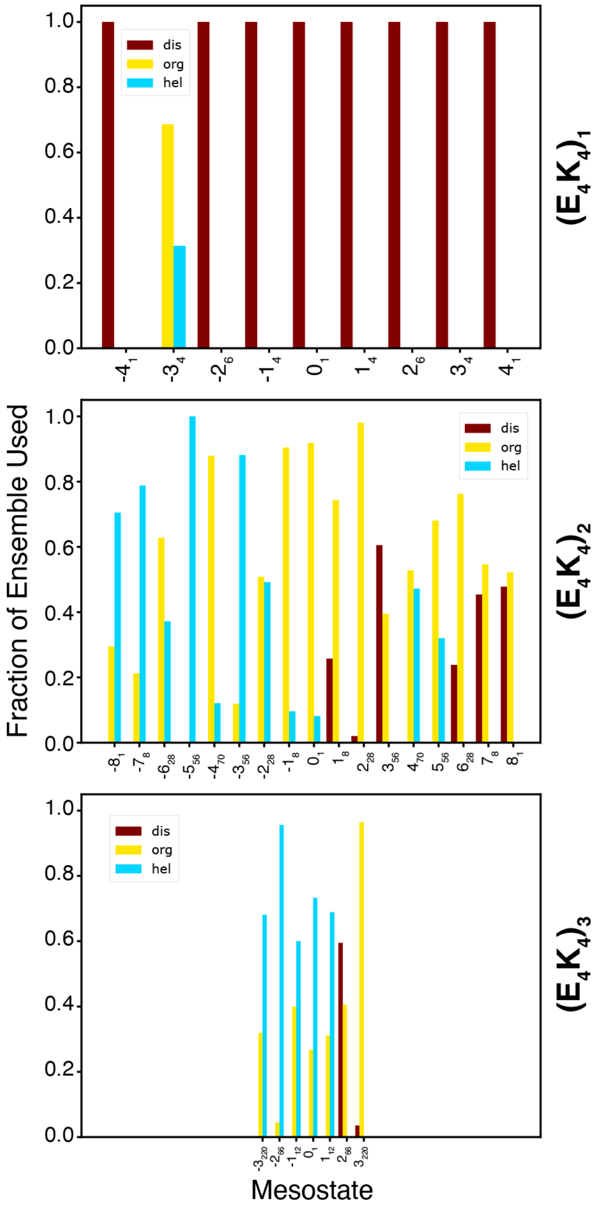


**Figure S6** : **Mesostate-specific weights for the mixing function used to derive pH-dependent probabilities for microstates using simulations of the disordered (dis), original (org) unmodified ensemble, and helical (hel) ensembles.** The weights that yield optimal agreements with the experimental data are summarized as mesostate-specific values The procedure used to arrive at the optimal combination of pH-dependent weights for mixing are summarized in **Figure 5**.

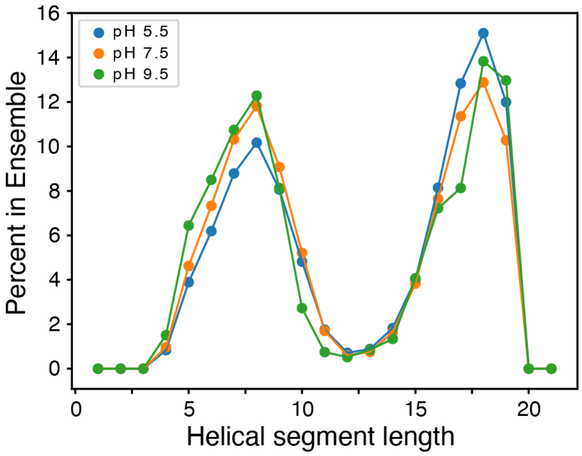


**Figure S7: Distributions of the lengths of helical segments extracted from the reweighted ensembles of $(E_4K_4)_2$ at pH 5.5, 7.5, and 9.5.**

### Supplementary references


(1) Kovács, Á.; Dudola, D.; Nyitray, L.; Tóth, G.; Nagy, Z.; Gáspári, Z. Detection of Single Alpha-Helices in Large Protein Sequence Sets Using Hardware Acceleration. *Journal of Structural Biology* **2018**, *204* (1), 109–116. https://doi.org/10.1016/j.jsb.2018.06.005.

(2) Gáspári, Z.; Süveges, D.; Perczel, A.; Nyitray, L.; Tóth, G. Charged Single Alpha-Helices in Proteomes Revealed by a Consensus Prediction Approach. *Biochimica et Biophysica Acta - Proteins and Proteomics* **2012**, *1824* (4), 637–646. https://doi.org/10.1016/j.bbapap.2012.01.012.

(3) Süveges, D.; Gáspári, Z.; Tóth, G.; Nyitray, L. Charged Single α-Helix: A Versatile Protein Structural Motif. *Proteins: Structure, Function and Bioinformatics* **2009**, *74* (4), 905–916. https://doi.org/10.1002/prot.22183.

(4) Fossat, M. J.; Posey, A. E.; Pappu, R. V. Quantifying Charge State Heterogeneity for Proteins with Multiple Ionizable Residues. *Biophysical Journal* **2021**, *120* (24), 5438–5453. https://doi.org/10.1016/j.bpj.2021.11.2886.
